# Hierarchical Neural Circuit Theory of Normalization and Inter-areal Communication

**DOI:** 10.1101/2025.07.15.664935

**Authors:** Asit Pal, Shivang Rawat, David J. Heeger, Stefano Martiniani

**Affiliations:** Simons Center for Computational Physical Chemistry, Dept. of Chemistry, New York University, NY, USA; Center for Soft Matter Research, Dept. of Physics, New York University, NY, USA; Courant Institute of Mathematical Sciences, New York University, NY, USA; Dept. of Psychology, New York University, NY, USA; Center for Neural Science, New York University, NY, USA

## Abstract

The primate brain exhibits a hierarchical, modular architecture with conserved microcircuits executing canonical computations across reciprocally connected cortical areas. We present a hierarchical neural circuit theory with feedback connections that dynamically implements divisive normalization across its hierarchy. In a two-stage instantiation (V1 ↔ V2), increasing feedback from V2 to V1 amplifies responses in both areas. We analytically derive power spectra (V1) and coherence spectra (V1-V2) as functions of frequency (f), and compare them with experimental observations: peaks in both spectra shift to higher frequencies with increased stimulus contrast, and power decays as 1/f^4^ at high frequencies. The closed-form spectra are validated against direct stochastic simulation of the full nonlinear circuit across contrasts. The theory further predicts distinctive spectral signatures of feedback and input gain modulation. Crucially, the theory offers a unified view of inter-areal communication, defined as the ability to linearly predict neural activity in one brain area from neural activity in another, with emergent features consistent with empirical observations of both communication subspaces and inter-areal coherence. It admits a low-dimensional communication subspace, where inter-areal communication is lower-dimensional than within-area communication. It further predicts that: i) increasing feedback strength enhances inter-areal communication and diminishes within-area communication, without altering the subspace dimensionality; ii) communication is strongest, and the subspace dimensionality lowest, at the frequencies at which V1-V2 coherence peaks; iii) normalization reduces the subspace dimensionality. Finally, a three-area (V1 ↔ V4 and V1 ↔ V5) instantiation of the theory demonstrates that differential feedback from higher to lower cortical areas dictates their dynamic functional connectivity. Altogether, our theory provides a robust and analytically tractable framework for generating experimentally-testable predictions about normalization, inter-areal communication, and functional connectivity.

---

The primate brain is a complex hierarchical network of interconnected cortical areas that perform computations essential for sensorimotor processing and cognitive functions. Among these, certain computations are repeated across cortical areas and are thus said to be “canonical computations”. Divisive normalization [1] is one such fundamental canonical principle, playing a central role in regulating neural responses across sensory and higher cortical areas.

Feedback connections are ubiquitous throughout the primate brain, yet their precise computational role remains poorly understood. A wide range of functions has been attributed to feedback, including attentional modulation [2], inter-areal communication [3], learning [4], and generative modeling [5]. While divisive normalization has been extensively studied in feedforward recurrent networks [6–9], its implementation and effects in hierarchical networks with feedback remain unexplored.

Effective communication between brain areas is essential for complex cognitive tasks, yet the precise mechanisms of communication remain an active area of investigation. One prominent hypothesis, *communication through coherence* (CTC) [10], proposes that synchronized oscillations (coherence) are crucial for effective inter-areal communication. Indeed, coherence between brain regions has been widely observed across cognitive, perceptual, and behavioral tasks [11, 12]. However, the causal nature of the relationship remains a key unresolved question. It is currently debated whether coherence is a fundamental mechanism enabling inter-areal communication or rather an epiphenomenon emerging from underlying interactions [13, 14]. An alternative hypothesis suggests that brain areas communicate through a low-dimensional *communication subspace* (CS) embedded within the high-dimensional neural activity space [15]. This concept, also identified in artificial neural networks [16], posits that only activity within these shared subspaces between the source and target areas facilitates communication. Recent modeling work has begun to examine how communication subspaces and the directionality of inter-areal interactions arise from circuit dynamics and connectivity [17–20]. Despite strong empirical evidence for both coherence-based and subspace-based communication, a unified, analytically tractable framework that jointly accounts for coherence and communication subspaces and explaining how they are dynamically modulated by factors such as feedback and stimulus contrast is currently lacking.

To address these gaps, we introduce a hierarchical recurrent neural circuit theory (Fig. 1) built upon two *a priori* assumptions: i) a hierarchy of brain areas with feedforward, recurrent, and feedback connections, and ii) recurrently implemented normalization within each brain area. A concrete model also requires choices about the connectivity, the normalization pool, and the noise (Methods); their influence on the results is examined in Suppl. Fig. S4 and in the *SI Appendix* (“Robustness with respect to model parameters”). Within this framework, we derive analytical expressions for inter-areal coherence and communication subspaces, offering experimentally testable predictions (see Discussion, “Experimental predictions of the model”). Given these choices, and a single set of parameters, the theory gives rise to a rich set of emergent properties that account for a wide spectrum of neurophysiological phenomena, including modulation of contrast response functions, oscillatory dynamics evident in power and coherence spectra, the emergence of communication subspaces, and the dynamic control of functional connectivity. A central contribution of this work is a unified view of interareal communication. The theory simultaneously predicts empirical observations of both communication subspaces and inter-areal coherence, and further reveals that coherence actively shapes the communication subspace. In doing so, our framework reconciles two hypotheses that have hitherto been treated as alternatives, and shows that coherence and subspace structure are two emergent properties of the same underlying circuit.

Finally, we emphasize that throughout our analysis, we neither adjust model parameters (such as time constants, weight matrices, and noise amplitudes) to reproduce specific experimental trends nor fit data when making direct comparisons. All results presented were obtained using a *single* set of model parameters. The only exception is Suppl. Fig. S7, in which the parameters are fitted to measurements of LFP power spectra. Additionally, we have verified that the key results hold across a wide range of parameters; for further details, see the *SI Appendix*.

**Figure 1:**
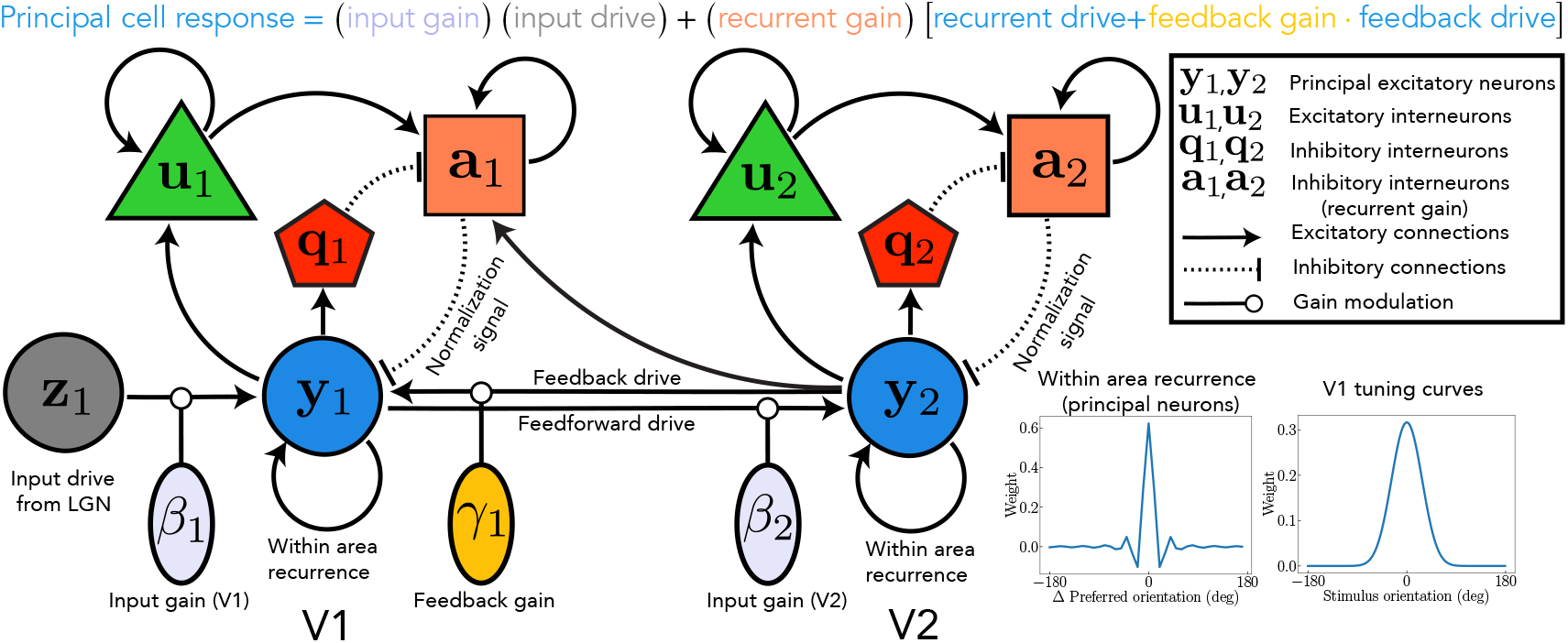
Schematic of the two-stage recurrent circuit model. Solid and dashed lines represent net excitatory and net inhibitory (effective) connections, respectively. Principal neurons are denoted by **y**, modulatory excitatory neurons by **u**, and modulatory inhibitory neurons by **a** and **q**. Subscripts 1 and 2 designate neurons in visual areas V1 and V2, respectively. **z**_1_ denotes the input drive from the LGN to V1, a weighted sum of the responses of LGN neurons. Parameters β_1_ and γ_1_ modulate the input and feedback gain to V1, respectively.

## Results

### Theory

Initially conceived to explain the responses in the primary visual cortex (V1) [1], the normalization model has emerged as a canonical computation across sensory and cognitive domains [21], including higher level vision, olfaction, audition, somatosensation, attention, multi-sensory integration, working memory, and decision-making. For citations see [6]. At its core, the normalization model proposes that the response of an individual neuron is scaled by the collective activity of its neighboring neurons (viz., the normalization pool), analogous to normalizing the length of a vector.

In addition, normalization decorrelates population responses to reduce redundancy [22, 23],maintains network stability in spite of recurrent amplification [8, 9], and generates long timescales without fine-tuning [24]. It is a computational-level description of neural responses that can be implemented using various mechanisms. Experimental evidence suggests that normalization operates via recurrent amplification [25, 26], amplifying weak inputs more than strong inputs, consonant with evidence that cortical circuits are dominated by recurrent excitation [27, 28].

**O**scillatory **R**ecurrent **G**ated **N**eural **I**ntegrator **C**ircuits (ORGaNICs) is a theoretical framework that dynamically implements divisive normalization by modulating recurrent amplification [6, 7]. Previous work on normalization models (including previous work on ORGaNICs) has focused on a single neural circuit within a brain region, such as V1, ignoring the abundant top-down feedback connections. Here, we develop a new hierarchical ORGaNICs theory with feedback connections in addition to feedforward and recurrent connections.

The hierarchical ORGaNICs theory describes neuronal population responses as dynamic processes evolving in a recurrent circuit, modeled by nonlinear differential equations (see Fig. 1 for illustration and Methods for equations). The circuit comprises distinct cell types: principal cells (**y**), excitatory modulator cells (**u**), and inhibitory modulator cells (**a, q**) (n.b., “modulator” refers to a multiplicative computation, not necessarily implemented with neuromodulators). Output firing rates of principal cells depend on three terms: 1) feedforward input drive modulated (viz., scaled) by input gain; 2) feedback drive modulated by feedback gain; 3) the sum of recurrent drive and scaled feedback drive, collectively modulated by recurrent gain (see equation at the top of Fig. 1). Whereas in Wilson–Cowan-type excitatory–inhibitory models apply the same form of rate equation to each subpopulation of neurons, with different signs and weights, the four populations here obey structurally different equations, each implementing a distinct operation (Methods, Eq. 1). Specifically, principal cells **y** sum the gain-modulated input, recurrent, and feedback drives (with a squaring output nonlinearity); excitatory modulator cells **u** accumulate the normalization signal, a weighted sum of principal-cell responses (square-root output nonlinearity); inhibitory modulator cells **a** implement the division, i.e., the multiplicative gain control of the recurrent and feedback drives; and inhibitory cells **q** track principal-cell activity and provide the sign inversion required by the feedback term in the dynamics of **a** (*SI Appendix*, Eqs. S21 and S22). This theory is particularly well-suited to analytical treatment, e.g., to derive closed-form solutions for the power spectral density, coherence, and communication subspaces (see Methods and Appendix).

Each neuron receives *input drive* defined as a weighted sum of responses from neurons at lower stages of the hierarchy. For example, LGN responses provide input drive to V1 (**z**_**1**_ in Fig.1), and V1 responses similarly drive V2 (solid arrow from **y**_**1**_ to **y**_**2**_ in Fig.1, representing principal neuron responses in each area). This weighting is specified by an embedding matrix of effective weights, which determine each neuron’s stimulus preferences, such as orientation selectivity (Fig.1, V1 tuning curves), analogous to a standard receptive field model (see Methods). Throughout, the weight matrices describe effective connectivity rather than anatomical synapses: negative weights may, for example, be implemented with disynaptic inhibition through interneurons that are not modeled explicitly (see Discussion). Input gain is determined independently by input modulator cells (*β*_1_ and *β*_2_ in Fig.1), whose gain may, in principle, depend on both afferent input and local population activity (e.g., responses of both LGN cells and V1 cells may contribute to the input gain in V1).

Each neuron also receives *recurrent drive* arising from weighted interactions among principal cells in the same brain area (Fig. 1, curved arrows). Recurrent connectivity follows a center-surround architecture, with local excitatory and distal inhibitory connections (Fig. 1, recurrent synaptic weights). The gain of this recurrent drive is dynamically controlled by recurrent modulator cells (**a**_1_ and **a**_2_), which combine excitatory (**u**_1_, **u**_2_) and inhibitory signals (**q**_1_, **q**_2_) from local interneuron populations as well as feedback from higher cortical areas (solid arrow from **y**_**2**_ to **a**_1_ in Fig. 1). Divisive normalization is achieved dynamically through the activity of these modulator cells, which carry the “normalization signal”.

Finally, each area receives *feedback drive*, a weighted sum of responses from neurons at higher stages of the hierarchy (e.g., V2 → V1; Fig. 1). The feedback drive is specified by a feedback weight matrix, which, for simplicity, we take to be the transpose of the corresponding feedforward matrix from V1 to V2 [5]. The strength of this feedback is controlled by a feedback gain parameter (*γ*_1_ in Fig. 1). More generally, feedback strength may be modulated either by changing the feedback gain or by changing the gain of the neural responses in the higher cortical areas providing feedback (see Discussion). Thus, the feedback gain may be a fixed property of the circuit or a dynamically regulated quantity that depends on both the inputs and outputs of the corresponding area. Importantly, even if the feedback gain is fixed, it may still be possible to manipulate the feedback strength experimentally to test predictions of the theory (elaborated in what follows).

The two-stage recurrent model with feedback implements divisive normalization simultaneously across multiple cortical areas. The implementation is exact in a special case: when the gains are set to *γ* = *β* = 1, and the recurrent weight matrix is the identity matrix (i.e., each neuron recurrently amplifies only itself), the steady-state responses of the model coincide precisely with the analytical steadystate solutions of the normalization equation (Methods, Eq. 2). For the center-surround recurrent weight matrix used in all results reported here, the circuit very closely approximates normalization (see Methods and Ref. [9]). The exact fixed point is obtained numerically by integrating the deterministic dynamics to steady state, and all spectra and subspaces are then computed in closed form by linearizing about this fixed point (see Methods).

#### The core mechanism in brief

Noise-driven fluctuations around the normalized fixed point are characterized by the cross-spectral density matrix ***S***(*ω*) (Methods, Eq. 6). Its diagonal entries determine the power spectra of individual neurons, while its normalized off-diagonal entries determine inter-areal coherence (Eq. 7). Integrating ***S*** (*ω*) over frequency yields the covariance matrix, and its frequency-resolved counterpart provides the covariance structure from which the communication subspace is obtained by reduced-rank regression (Methods). Thus, power, coherence, and communication-subspace structure are complementary readouts of the same underlying fluctuation dynamics.

Recurrent normalization shapes these dynamics in two related ways. First, recurrent gain control modifies the circuit dynamics around the fixed point, giving rise to contrast-dependent resonances that appear as peaks in both power and coherence. Second, competition within the normalization pool preferentially suppresses weakly driven neurons, producing a soft winner-take-all effect that reduces their contribution to the cross-covariance and thereby lowers the effective dimensionality of the communication subspace, particularly near the resonant frequencies. Removing normalization abolishes the resonant peaks in power and coherence and produces a higher-dimensional communication subspace (Suppl. Figs. S4e–h and S5).

### Contrast response functions

Both V1 (dashed) and V2 (solid) exhibit the characteristic saturating, sigmoidal dependence on contrast (As shown in Fig. 2a, b). The model predicts a lower semisaturation constant (the contrast at which the firing rate reaches half-saturation) and a steeper slope for cortical area V2 compared to V1. This prediction is commensurate with experimental observations in the literature that used optimal stimulus sizes (Fig. 2b) [29, 30]. Similar hierarchical trends, specifically an increase in slope and a decrease in the semisaturation constant, have also been observed across other brain areas like the LGN, V1, and MT with progression up the visual hierarchy [32].

**Figure 2:**
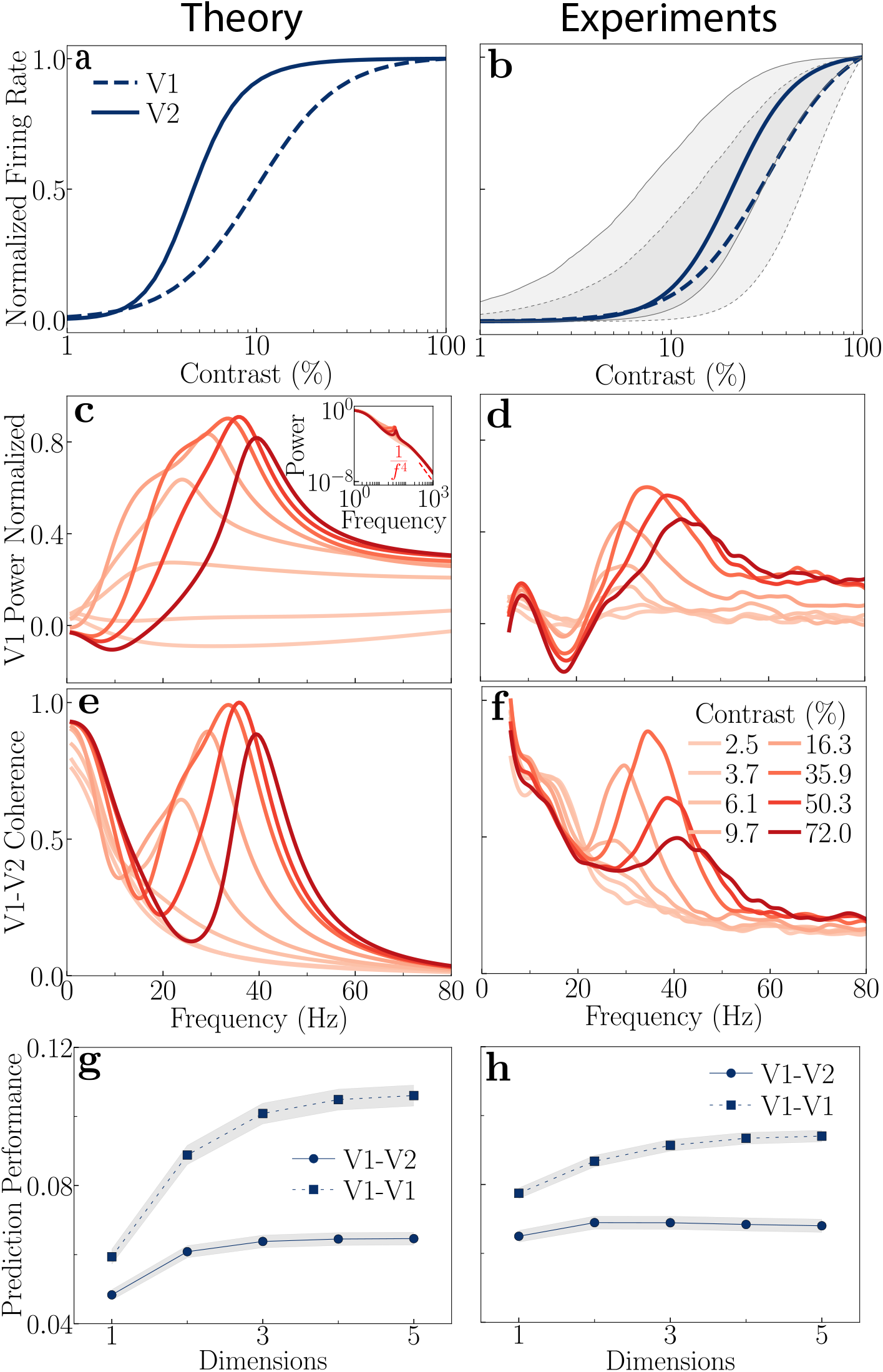
Comparison of theoretical predictions (left column) and experimental observations (right column) in V1 and V2. All theoretical predictions are generated using the same baseline parameters (Table S1). The analytical spectra in **c** and **e** are validated against direct stochastic simulation of the full nonlinear circuit in Suppl. Fig. S6, and a quantitative comparison in which the model parameters are fitted to experimental measurements of the power and coherence spectra is given in Suppl. Fig. S7. **a**, Theoretical mean firing rates as a function of stimulus contrast (V1: dashed line, V2: solid line). **b**, Experimental mean firing rates of V1 and V2 (data from [29] and [30], replotted for comparison). The shaded area with a solid border indicates the 25th to 75th percentile range for V1, and the one with the dashed border indicates the same for V2. **c**, Theoretical V1 power spectra at various stimulus contrast levels. Power spectra were normalized using the equation 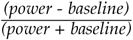, where baseline power is taken to be spontaneous activity at 0% contrast. The peak power shifts towards higher frequency with increasing contrast. The inset shows the 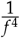 power law decay at high frequency. **d**, Experimental power spectra from macaque V1 (data from [31], replotted for comparison). **e**, Theoretical coherence spectra (normalized) between maximally firing neurons in V1 and V2. The peak coherence shifts towards higher frequency with increasing contrast. **f**, Experimental coherence spectra from macaque V1-V2 (data from [31], replotted for comparison). **g**, Theoretical communication subspaces, prediction performance as a function of dimensionality for inter-area (circles) and within-area (squares) communication. Simulation results indicate that the inter-area communication subspace is lower dimensional than the within-area communication. **h**, Experimental communication subspaces from macaque V1-V2 (data from [15], replotted for comparison). Th**5**e plotted prediction performance in both panels **g** and **h** is an average across different subsets of the source and target neural populations, and the shaded areas represent the standard error of the mean (SEM).

Increasing either the feedback gain, *γ*_1_, or the V1 input gain, *β*_1_ (with the V2 input gain *β*_2_ held at its baseline value), enhances neural responses across cortical areas (Figs. 3a,b). Importantly, this response enhancement is evident primarily as a change in contrast gain, i.e., a shift of the contrast-response functions on the log contrast axis rather than a change in response amplitude. Increasing the V2 input gain, *β*_2_, leaves V1 responses unchanged and enhances only V2 responses (Suppl. Fig. S8c).

**Figure 3:**
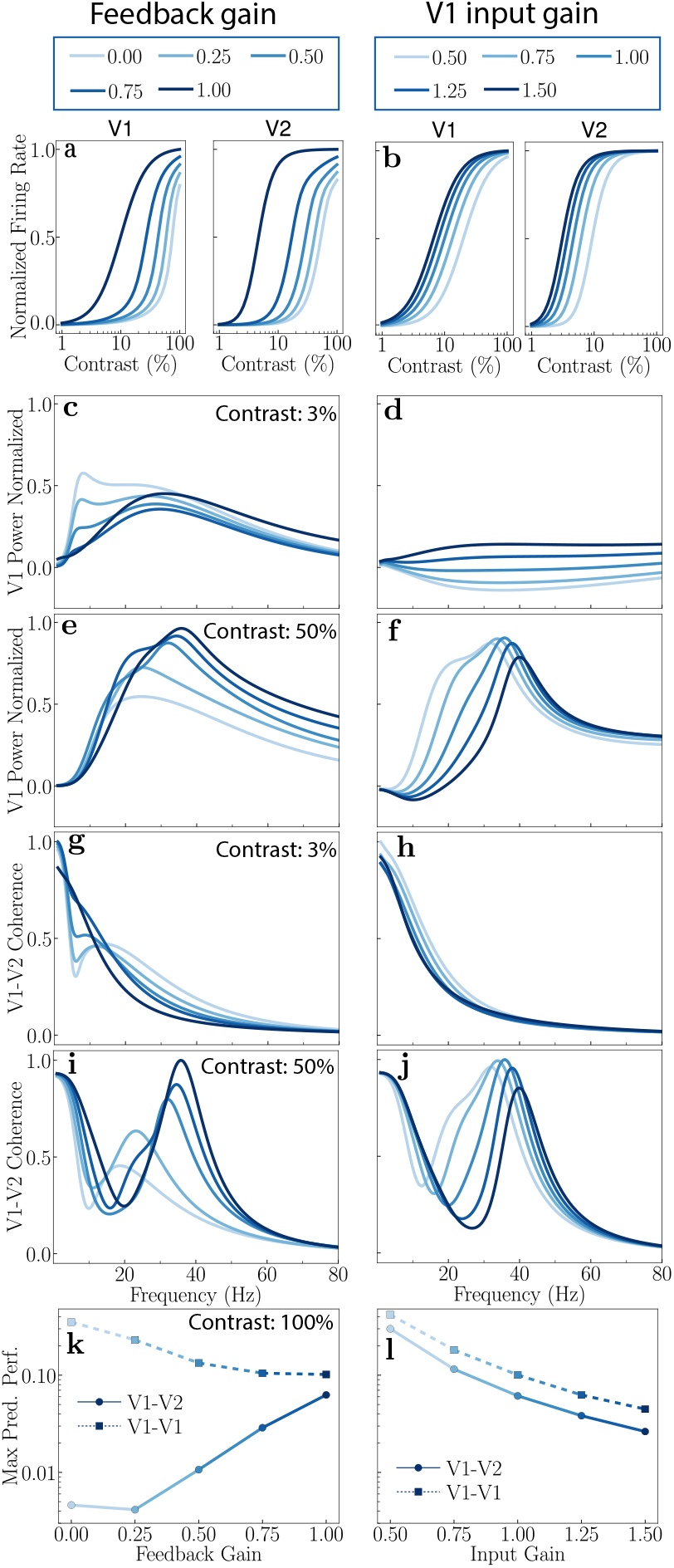
Theoretical predictions: Modulating feedback (left) and V1 input gain (right). **a**,**b**, Firing rates as a function of contrast. Increasing either feedback gain or V1 input gain enhances neural responses, with feedback gain showing a greater impact in the higher cortical area V2. **c**,**d**, V1 power spectra: 3% contrast. An alpha peak is observed for low feedback gain, but the peak diminishes with increasing feedback gain. The alpha peak is absent with V1 input gain changes. **e**,**f**, V1 power spectra: 50% contrast. A consistent gamma peak is observed, which shifts toward higher frequencies with increasing feedback gain (**e**) and V1 input gain (**f**). **g**,**h**, Coherence spectra: 3% contrast. A broad peak in the beta band is observed at low feedback gain, which vanishes at high feedback gain. No such peak is observed with changes in V1 input gain. **i**,**j**, Coherence spectra: 50% contrast. A beta peak is observed for low feedback gain, which shifts toward higher (gamma) frequencies with increasing feedback gain. No beta peak is observed for changes in V1 input gain, but the gamma peak shifts toward higher frequencies with increasing V1 input gain. **k**,**l**, Communication subspaces. Increasing feedback gain enhances inter-areal (circles) communication while decreasing within-area (squares) communication. Conversely, increasing V1 input gain decreases both inter- and within-area communication. The V2 input gain β_2_ is held at its baseline value (1.0) throughout; the effect of varying β_2_ is shown in Suppl. Fig. S8.

### V1 power spectra

Power spectrum analysis of electroencephalographic (EEG) and local field potential (LFP) signals has been used extensively to identify and quantify oscillatory activity in various frequency bands (e.g., alpha, beta, gamma, theta), hypothesized to be linked to various cognitive functions, behavioral states, and neurological conditions [33]. Since our model has a known fixed point solution, we calculate the power and coherence spectra analytically, given a model of the noise in the circuit (see Methods). The analytical spectra rest on a linearization of the dynamics about the fixed point; we verified them against direct numerical integration of the full nonlinear stochastic circuit at every contrast (Suppl. Fig. S6).

We adopted an analytically tractable noise model that adds low-pass filtered synaptic noise to the membrane potential and multiplicative Gaussian white noise to the firing rate dynamics. Furthermore, we explicitly accounted for long-range correlated noise in LFP measurements. This type of noise, propagating through brain tissue, affects LFPs with negligible impact on spiking activity and firing rates [34]. Further details are provided in the *SI Appendix* (“Intrinsic neural circuit noise” and “Extrinsic noise in the local field potential”).

Our theory reproduces the main qualitative features of the V1 power spectra across varying contrast levels. These theoretical V1 predictions (Fig. 2c) are compared with experimental spectra from macaque V1 (Fig. 2d) under identical contrast conditions [31]. With the unfitted baseline parameters, the model captures three qualitative trends: 1) the power spectrum of the noise is shaped by recurrent normalization to exhibit a resonant peak in the gamma-frequency range (≈ 40-60 Hz); 2) the peak frequency shifts right-ward with increasing contrast; and 3) the peak power decreases after reaching a maximum around 40% contrast. A quantitative comparison, in which the model parameters are fitted to the measured spectra, is provided in Suppl. Fig. S7a (*SI Appendix*, “Quantitative fit to experimental power and coherence spectra”). The oscillatory activity in the gamma-frequency range depends on normalization; removing normalization eliminates the peaks (see Suppl. Fig. S4e), commensurate with experimental observations that gamma oscillations are linked to normalization [35, 36]. The peak frequencies depend on the values of the time constants; larger time constants would shift the peaks to lower frequencies (see Discussion: Limitations).

The theory also reproduces the steep fall-off in power reported at higher frequencies [37], where power approximately scales as 1/ *f* ^4^ with increasing frequency (*f*) (Fig. 2c, inset). This occurs because the synaptic noise added to the membrane potentials is first low-pass filtered by the synapse’s time constant, and then by recurrent processing in the circuit, which also acts as a low-pass filter (each contributing a factor of 1/ *f* ^2^ to the power spectrum).

The theory offers a novel, experimentally testable prediction that modulating feedback, feedforward, and input gains elicits distinct changes in the power spectra. Figures 3c-f illustrate these distinct effects on the normalized power spectra when feedback gain (left panels) or input gain (right panels) is varied. These comparisons are presented for both a low (3%; Figs. 3c,d) and a high (50%; Figs. 3e,f) stimulus contrast. See Supplementary Material for comprehensive results across other contrast levels (Suppl. Fig. S1), and Suppl. Fig. S8d–i for the effect of modulating the V2 input gain.

The model predicts oscillatory activity in the alpha-frequency range (≈ 10 Hz) for low stimulus contrasts and small feedback gain values (Fig. 3*c*). As the feedback gain to the principal cells increases, this alpha peak diminishes and ultimately vanishes at high feedback gain levels, while a broad peak concurrently emerges in the gamma frequency range (Figs. 3c,e). No such oscillatory activity in the alpha-frequency range is observed by manipulating the V1 input gain, for any contrast (Figs. 3d,f).

The power spectra are dominated by oscillatory activity in the gamma-frequency range at high stimulus contrast, regardless of the V1 input gain and feedback gain values (Figs. 3e,f). Increasing the feedback gain causes this gamma peak to shift towards higher frequencies. A similar shift of the gamma peak to higher frequencies is observed when the V1 input gain is increased. However, a key distinction arises in their impact on the peak’s bandwidth: the gamma peak bandwidth narrows with increasing V1 input gain but broadens with increasing feedback gain.

### Inter-areal coherence

Coherence measures the statistical correlation between two signals (e.g., LFPs in two different brain areas) as a function of frequency. Neuronal synchrony is hypothesized to contribute to the dynamic selection and routing of information both within and between brain areas [10, 38, 39].

Our model’s predictions for V1-V2 coherence spectra accurately capture key experimental trends observed across various stimulus contrasts. We analytically determined the coherence between the maximally firing neurons of V1 and V2 (see Methods). Figure 2e displays the theoretically predicted V1-V2 coherence spectra for different contrasts. These model predictions are compared with experimental V1-V2 coherence data from macaque monkeys, obtained under identical contrast conditions (Fig. 2f; [31]). The model accurately captures three qualitative observations: 1) high coherence at lower frequencies for low contrasts; 2) a shift of the peak coherence toward higher frequencies with increasing contrast; and 3) a decrease in peak coherence after reaching a maximum around 40% contrast. The coherence peaks in the gamma frequency range are due to normalization; removing normalization eliminates the peaks in coherence (see Suppl. Fig. S4f). Suppl. Fig. S7b shows model fits to experimental measurements of the coherence spectra. Feedback and input gain differentially modulate V1-V2 coherence across contrasts (Figs. 3g-j; Suppl. Fig. S8j–o). At low contrast (3%), a coherence peak in the beta band (12-30 Hz) is observed with low feedback gain (Fig. 3g); this peak disappears as feedback gain is increased. Notably, at low feedback gain, a dip in V1-V2 coherence occurs in the alpha band, corresponding to the frequencies where peak power is observed in V1. This suggests that alpha band activity, despite its prominence in V1 power, may not effectively propagate to V2 or participate in inter-areal communication. No such peaks are observed with changes in the V1 input gain for low contrast (Fig. 3h). At high contrast (50%), increasing feedback gain shifts the peak coherence towards higher frequencies and increases both its magnitude and width (Fig. 3i). A similar frequency shift in peak coherence occurs with increasing V1 input gain, but its magnitude and width decrease (Fig. 3j). Increasing the V2 input gain shifts the coherence peak to higher frequencies, broadens it, and reduces its magnitude, and produces no beta-band peak at low contrast (Suppl. Fig. S8l,o).

### Communication subspaces

Neural communication is hypothesized to be channeled through “communication subspaces (CS)” [15]. Although neural activity space is high-dimensional, neural communication may not utilize this vast space. CS are lower-dimensional manifolds within the broader neural activity space that preferentially engage in inter-areal communication. In contrast, within-area communication utilizes a distinct, higher-dimensional “private” subspace [15].

Our theory successfully captures key characteristics of the CS (Figs. 2g,h). We derived an analytical model to examine both inter-areal (V1-V2) and within-area (V1-V1) communication (see Methods). We found that the CS is low-dimensional, much lower than the number of sampled neurons (source and target populations of 30 neurons in Figs. 2g and 3k,l; see Methods); the dependence of the dimensionality on the number of modeled and sampled neurons is characterized in Suppl. Fig. S9. In addition, the inter-areal communication (V1-V2) subspace is lower-dimensional than the within-area (V1-V1) communication subspace. Figure 2g illustrates the model prediction performance (i.e., the ability to predict activity fluctuations of a target subpopulation from the activity fluctuations of a source subpopulation using a linear model) as a function of the dimensionality of the CS. These theoretical results are qualitatively consistent with the experimental data (Fig. 2h; [15]).

Feedback and input gain differentially modulate communication subspaces (Figs. 3k,l; Suppl. Fig. S8p– r). Increasing feedback gain from the higher cortical area (V2) to the lower cortical area (V1) enhanced inter-areal communication while diminishing within-area communication (Fig. 3k), without substantially altering the subspace dimensionality (see Suppl. Figs. S3a,b). Conversely, increasing V1 input gain reduced both the inter-area and within-area communication (Fig. 3l). These results are for 100% contrast; see Suppl. Fig. S3c for other contrasts. Increasing the V2 input gain, *β*_2_, enhances inter-areal communication, like feedback gain, but leaves within-area communication approximately unchanged (Suppl. Fig. S8r). However, the effect of feedback gain is substantially larger than that of V2 input gain. The observed enhancement of inter-areal communication with increased feedback suggests that gain modulation, and top-down feedback in particular, can dynamically modulate the efficiency of information transfer between cortical areas without altering the underlying structure of the communication subspace (see below, Dynamic modulation of functional connectivity through feedback), a finding consistent with previous work on subspace dimensionality [40].

### Frequency decomposition of communication subspaces

The theory predicts that the effectiveness of communication between brain areas varies systematically with frequency and contrast (Fig. 4). We quantified the efficacy of communication by analytically calculating the prediction performance for each frequency component (see Methods). Using the Wiener-Khinchin theorem, we converted the power spectra of the source and target population into their noise correlations. To isolate noise correlations at a specific frequency, we applied an idealized narrow-band pass filter. This conceptual filter is implemented via a transfer function, which is non-zero only within a very narrow frequency band around the frequency of interest. Applying this filter to the power spectra and integrating across the frequency domain yields the frequency-specific correlations (see Methods, “Frequency decomposition of the communication subspace”. This conversion enabled us to predict the power of the target population from the source population (using a linear readout) at each frequency, thus providing a measure of communication efficacy (prediction performance) as a function of frequency.

**Figure 4:**
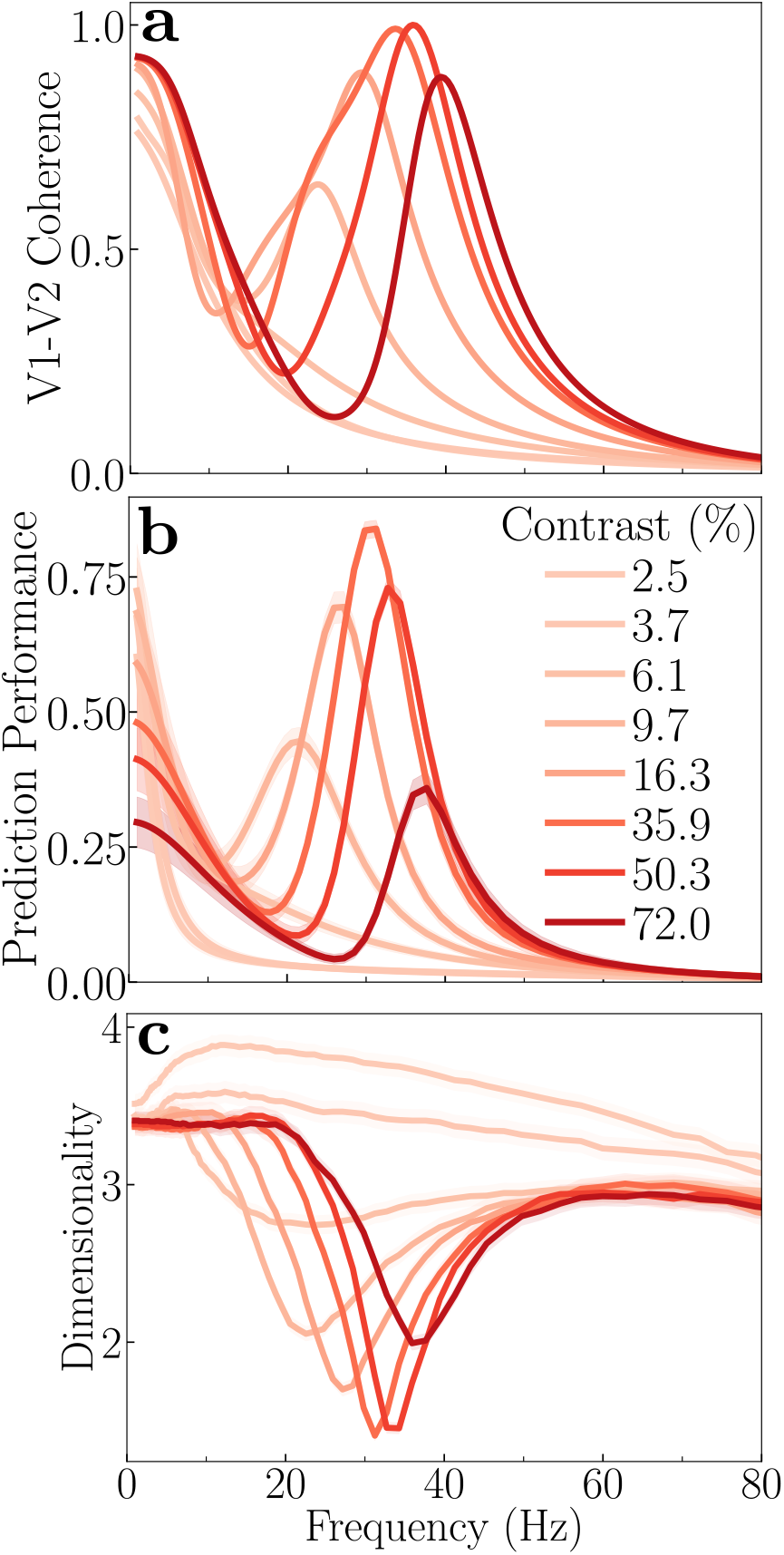
Theoretical prediction: Frequency-dependence of communication. **a**, V1-V2 coherence spectra for different contrasts; same as in Fig. 2e. **b**, Prediction performance versus frequency at different contrasts, averaged across different subsets of the source and target populations. At low stimulus contrast (≈ 10 %), a peak in communication efficacy is observed around 20 Hz. As stimulus contrast increases, the preferred frequency of communication shifts towards higher frequencies (40 Hz). The magnitude of communication reaches a maximum around 40% contrast, before declining at higher contrast levels. Under conditions of very low contrast (< 10%), communication is mostly concentrated at very low frequencies. These frequency-specific trends in communication parallel the V1-V2 coherence patterns shown in panel **a. c**, Dimensionality of the communication subspace versus frequency, averaged across different subsets of the source and target populations. A notable dip in dimensionality occurs at frequencies corresponding to peaks in communication (panel **b**) and coherence (panel **a**). The shaded areas in panels **b** and **c** represent the standard error of the mean (SEM) across different subsets of the source and target populations.

V1-V2 communication is dynamically modulated by stimulus contrast in a frequency-dependent manner. At low stimulus contrasts (e.g, approximately 10%), a distinct peak in communication is observed at ≈ 20 Hz (Fig. 4b). As stimulus contrast increased, this preferred frequency for communication shifts toward higher frequencies (Fig. 4b). Furthermore, the magnitude of this communication peak exhibits a non-monotonic relationship with contrast, reaching its maximum at around 40% contrast before declining at higher contrasts. These communication peaks are due to normalization; removing normalization eliminates the peaks in both coherence and prediction performance (see Suppl. Figs. S4f,g). Under conditions of very low contrast, no distinct peak in communication is observed; instead, communication is mostly concentrated at very low frequencies (< 10 Hz). These frequency-specific trends in communication mirror the V1-V2 coherence spectra (Fig. 4a). This correspondence is a direct consequence of how the two quantities are defined: coherence (Eq. 7) and frequency-resolved prediction performance (Eq. 22) are both computed from the same cross-spectral density matrix, so the frequencies at which the source and target populations are most strongly correlated are also those at which one best predicts the other.

The dimensionality of the CS is the number of dimensions of source activity required to reach the full-rank prediction performance (within a small tolerance); it is dictated by the rank of the covariance matrix between the source (V1) and target (V2) subpopulations (see Methods: Interpretation of the results, Suppl. Fig. S5). The dimensionality of the inter-areal (V1-V2) CS also varies with frequency and stimulus contrast (Fig. 4c). The dimensionality of the communication subspace decreases at frequencies where inter-areal coherence and communication are maximal, for each contrast (Fig. 4c).

The dimensionality of the CS is influenced mainly by three factors. First, dimensionality depends on normalization. Although dimensionality is relatively low in the absence of normalization (Suppl. Fig. S4h), including normalization further reduces dimensionality at coherent frequencies (Fig. 4c). The intuition for why normalization reduces dimensionality is that when only a few neurons are strongly driven, and their responses are highly correlated across areas (V1 and V2), normalization suppresses the activity of other nearby neurons that are not strongly driven. This suppressive interaction, akin to a soft winner-take-all mechanism, diminishes the contribution of other neurons to the shared covariance, thereby reducing the overall dimensionality. Second, the dimensionality is increased for a fully interconnected (all-to-all) connectivity between V1 and V2 (Suppl. Fig. S4l). Third, delocalized inputs (e.g., noise images or naturalistic images) lead to a higher dimensionality than localized inputs (e.g., gratings) (Suppl. Fig. S4p), commensurate with experimental results [15].

### Dynamic modulation of functional connectivity through feedback

Anatomical connections influence neural communication but do not completely determine it. Rather, it has been observed experimentally that functional connectivity between brain areas changes dynamically [41, 42].

Our theory predicts that functional connectivity is controlled by modulating feedback strength (Fig. 5). We expanded our model to encompass three cortical areas: V1, V4, and V5→ (also called MT). In this expanded model, V1 has reciprocal connections with both V4 and V5, representing the parallel dorsal and ventral visual pathways, respectively [43]. For simplicity, we assume there are no direct connections between V4 and V5 in the model. Analysis of this three-area model provides a demonstration that differential feedback strength can control functional connectivity. When the feedback strength from V5 → V1 is greater than the feedback from V4 → V1, we observe that V1 strengthens communication with V5 (Fig. 5a). Conversely, when the feedback from V4 to V1 is stronger, V1 shows enhanced communication with V4 (Fig. 5b). This suggests that the relative strength of feedback from the higher cortical area (V5 or V4) can dictate the information flow from the lower cortical area (V1). One possible way this could be achieved is by selectively enhancing the neural activity in either V5 or V4, which will, in turn, increase the feedback to V1. This is commensurate with experimental evidence that feature-based attention and task demands may selectively boost activity in one or another cortical area [44], and specifically that attending to motion increases the transfer of motion information (communication) between V1 and V5 [45]. Therefore, top-down signals can dynamically modulate functional connectivity from lower to higher cortical areas, simply by adjusting the gain of neural activity in one higher cortical area (e.g., V4) relative to another (e.g., V5). This mechanism enables flexible cortical information routing, which is essential for cognitive flexibility. Because increasing the V2 input gain also enhances inter-areal communication (Suppl. Fig. S8r), we expect that differential V2 input gain to V4 versus V5 would route communication in the same way although to a lesser extent; the two mechanisms are distinguished by their effect on within-area communication, which is reduced by feedback gain but unchanged by V2 input gain, and by their spectral signatures (Suppl. Fig. S8).

**Figure 5:**
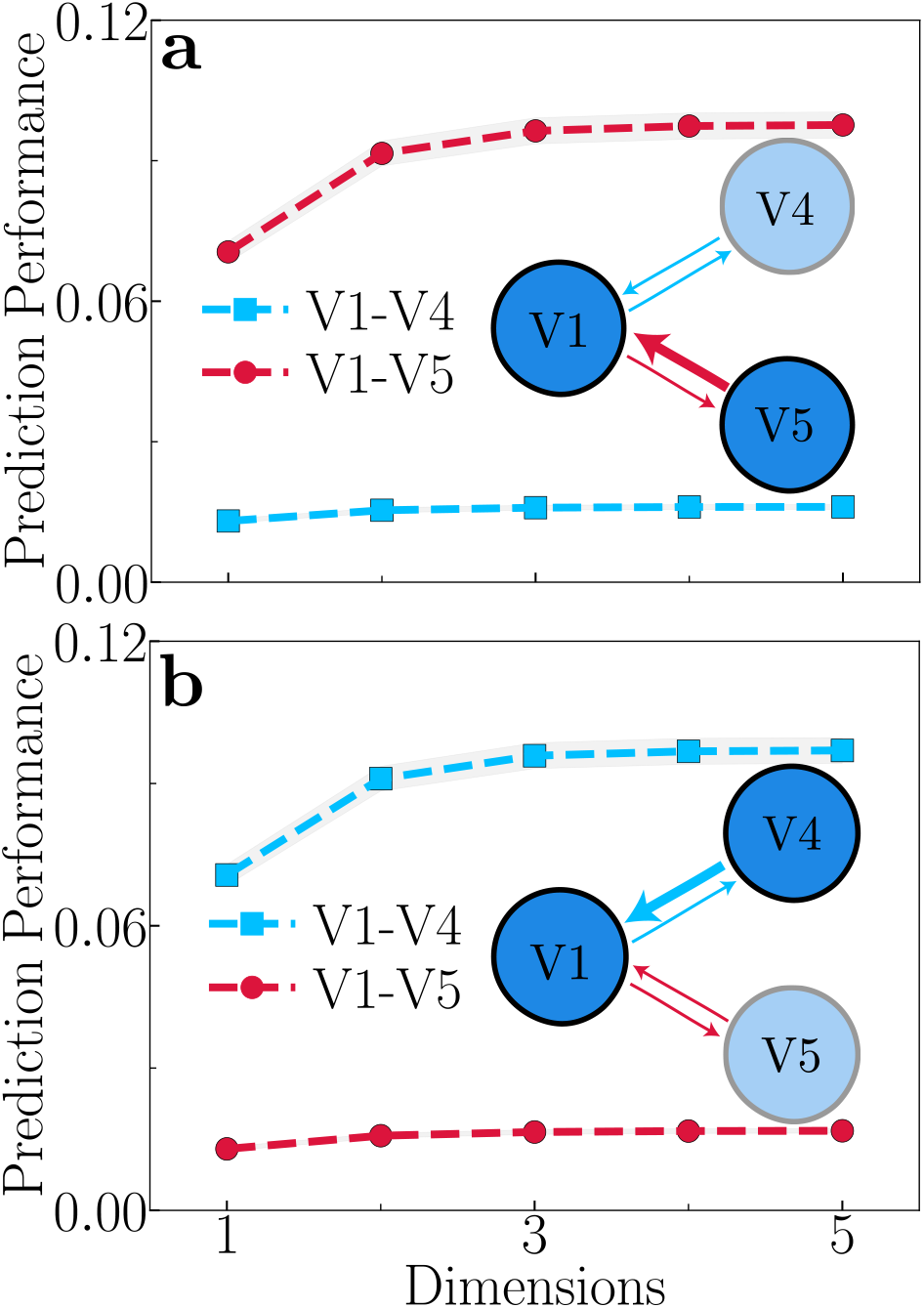
Theoretical prediction: Feedback-dependent modulation of functional connectivity. **a**, Feedback from V5 → V1 is stronger than feedback from V4 → V1, and V1 preferentially enhances communication with V5. **b**, Feedback from V4 → V1 is stronger than feedback from V5 → V1, and V1 preferentially enhances communication with V4. The plotted prediction performance is an average across different subsets of the source and target populations, and the shaded areas represent the standard error of the mean (SEM). These results demonstrate that top-down feedback can dynamically route information flow between cortical areas, i.e., modulating functional connectivity.

Note that, to isolate the variable of interest, we modeled V4 and V5 with identical parameters for their recurrent, feedforward (V1 → V4/V5) and feedback (V1 ← V4/V5) connectivity. This ensures that the observed increase in communication with either V4 or V5 is a direct consequence of the stronger feedback from that specific area.

## Discussion

We present an analytically tractable theory of neural circuit function that implements divisive normalization dynamically across interconnected brain areas. Emergent properties of the theory account for a diverse array of neurophysiological phenomena, including oscillatory activity (measurable as peaks in LFP power and coherence spectra) and communication subspaces. Based on the theory, we propose a unified view of inter-areal communication.

Our theory yields two key, experimentally-testable predictions. First, the dimensionality of the communication subspace is lowest at the frequencies at which V1-V2 coherence and communication peak, and these frequencies shift with contrast. Second, increasing feedback strength selectively enhances inter-areal communication and diminishes within-area communication, without altering the subspace dimensionality, whereas increasing the V2 input gain enhances inter-areal communication without reducing within-area communication. Candidate experimental manipulations and readouts for each of these predictions, and for the gain manipulations discussed below, are given in “Experimental predictions of the model”. A comprehensive list of further model results and their robustness is provided in *SI Appendix* section “Robustness with respect to model parameters”.

Furthermore, the theory proposes a mechanism for dynamic functional connectivity: top-down feedback signals enhance communication between brain areas simply by adjusting the gain of neural activity in one or more cortical areas. This modulation of neural gain offers a mechanism for dynamically changing the functional connections between brain areas over time.

### Neural circuit function

The original ORGaNICs model [6, 7], in its initial form without feedback connections, serves as a biophysically-plausible counterpart to Long Short Term Memory units (LSTMs) [46], a class of artificial recurrent neural networks (RNNs) prevalent in machine-learning applications [47]. Unlike LSTMs or any other previously proposed RNNs, ORGaNICs have built-in recurrent normalization, which offers several advantages, including stability even with exuberant recurrent connections [8, 9], and long timescales without fine-tuning [24].

The hierarchical version of the ORGaNICs theory introduced here adds feedback connections, drawing a partial analogy to “hierarchical predictive coding” models of sensory and perceptual processing [48–53]. However, our hierarchical theory diverges from predictive coding in its fundamental mechanisms. In predictive coding, it is proposed that forward-propagating signals represent prediction errors, while feedback pathways convey predictions to lower levels of hierarchy. The goal is to minimize prediction error by matching sensory input with top-down predictions. In hierarchical ORGaNICs, instead of propagating prediction errors, the circuit balances the input, recurrent and feedback drives of each principal cell (Eq. 1a) so that the steady-state responses follow the normalization equation (Eq. 2). This mechanism is more consistent with the observed physiology than predictive coding models. For example, neurons maintain sustained activity in response to a constant, fully predictable stimulus [54], whereas in predictive coding the forward-propagated activity, representing prediction error, should be “explained away” as the stimulus becomes predictable.

### Normalization

The proposed theory admits an analytical fixed point that follows the normalization equation (Eq. 2) *exactly*, for arbitrary non-negative normalization weights, when the input drive is constant over time, the recurrent weight matrix is the identity, and the input gain and feedback gain are equal to one (*β* = *γ* = 1) (see *SI Appendix* for derivation) [8, 9]. For the center-surround recurrent weights used here, the fixed point approximates normalization closely (Methods). Several circuit mechanisms have been hypothesized to underlie normalization, including shunting inhibition, synaptic depression, and inhibition-stabilized networks [26, 55, 56]In these models, normalization is realized either largely feedforward, through a divisive conductance or synaptic depression at the input, or approximately, as an emergent property of recurrent excitatory–inhibitory dynamics. The present circuit differs in three respects: the normalization equation, with an arbitrary normalization pool, is a fixed point of the recurrent dynamics rather than a description fitted to their steady state; the dynamics remain stable even with exuberant recurrent connections [8, 9]; and, because the fixed point and the linearized dynamics about it are available, the same circuit yields power spectra, coherence, and communication subspaces in closed form (Methods), which is what allows the spectral and population-level phenomena examined here to be traced to a single mechanism.

Our model achieves normalization dynamically via recurrent amplification, which is inversely regulated by the activity of inhibitory modulator cells. When the input is weak, the modulator cells exhibit a small response, leading to strong recurrent amplification. Conversely, a strong input generates a large response from modulator cells, which suppresses the amplification. This process modulates both excitatory and inhibitory recurrent signals, which is consistent with the experimental observation that normalization involves a decrease in both recurrent excitatory and inhibitory conductances [26]. Because the theory implements normalization, it is commensurate with a wide range of physiological and psychophysical phenomena previously explained by the normalization model (see Introduction for references).

The theory links normalization with oscillatory activity. Removing divisive normalization eliminates oscillatory peaks in power and coherence spectra (Suppl. Figs. S4e,f) and abolishes the link between coherence, prediction performance, and subspace dimensionality (Suppl. Figs. S4g,h), demonstrating that normalization is central to both the dynamics and function of the circuit.

### Role of feedback

This paper provides an experimentally testable computational model of cortical function where neural activity is shaped by a combination of feedforward drive (bottom-up) and feedback drive (top-down). Feedback in the model may serve a variety of functions. First, feedback provides a means for controlling inter-areal communication (functional connectivity) simply by changing the gain of neural activity in one or more cortical areas (see Discussion section “Attention and functional connectivity”). Second, the feedback drive may contribute to selective attention (see Section “Attention and Functional Connectivity”). Third, the relative strength of the input gain and feedback gain determines whether neural responses are driven bottom-up, top-down, or a combination of the two. Fourth, feedback contributes to the local normalization signal by providing excitatory drive to inhibitory neurons (see Methods). This feedback-mediated inhibition mechanism is essential for maintaining network stability, and is commensurate with experimental evidence that normalization depends in part on loops through higher visual cortical areas [57].

The interplay between the input gain and the feedback gain determines the primary driver of neural responses. If the input gain is strong and the feedback is weak, the system behaves like traditional feedforward models. Conversely, with strong feedback and weak input, the network is hypothesized to behave like generative models, creating sensory representations from higher-level abstract knowledge, akin to recalling an image from memory. When input and feedback are balanced, the framework may integrate prior beliefs with sensory evidence (see [5] for details), akin to models based on Bayesian inference [58, 59].

The theory predicts differential effects of changing input gain and feedback gain on the power, coherence spectra, and inter-areal communication (see Fig. 3 and Suppl. Fig. S8). Hence, these metrics may be used as a signature for distinguishing changes in input gain from changes in feedback (“Experimental predictions of the model”).

### Unified view of inter-areal communication

Our theoretical framework offers a novel perspective on inter-areal communication. Rather than viewing “communication through coherence” (CTC) and “communication subspace” (CS) as alternative hypotheses, our theory unifies them, showing that coherence and communication subspaces are complementary empirical phenomena emerging naturally from the same underlying mechanism. A similar conclusion was recently reached by Ni et al. [60] using a large-scale spiking neural circuit model. The frequency decomposition of the communication subspace mirrors the inter-areal coherence at every contrast. Communication is strongest at the frequencies of peak V1-V2 coherence, and the communication subspace is lowest-dimensional at the same frequencies. The reason is that coherence, prediction performance, and dimensionality are all computed from the same cross-spectral density matrix. Normalization shapes this matrix in two ways. First, normalization creates a resonance. This resonance sets the peaks of the coherence and of the prediction performance. Second, normalization suppresses weakly driven neurons. This suppression lowers the rank of the cross-spectral density matrix at the resonant frequencies and thereby reduces the dimensionality of the communication subspace.

### Attention and functional connectivity

The theory provides a possible explanation for the efficient routing of sensory information between cortical areas, i.e., for changing functional connectivity. Lower cortical areas, such as V1, can flexibly relay information to specific higher cortical areas (e.g., V4 vs. V5) based on the relative strength of the feedback from each of the higher cortical areas. The strength of the feedback can be controlled either by changing the feedback gain or (perhaps more simply) by changing the gain of the neural responses in each of the higher cortical areas. We hypothesize that this feedback-mediated process accounts for changes in functional connectivity [61] and feature-based attention [62]. Consistent with this hypothesis, experimental evidence demonstrates that feature-based attention is spatially global, i.e., that neural activity is boosted in feature-selective neurons with receptive fields covering the visual field [63].

The theory may also provide insights into the neural processes underlying spatially-selective attention with the addition of a retinotopic map. The theory identifies two distinct pathways through which attention modulates neural gain. First, attention may modulate the input gain of the sensory neurons, consistent with the established normalization model of attention [64]. Second, attention may modulate the activity of the sensory neurons via the top-down feedback drive. Exogenous attention is the fast, involuntary orienting of attention driven by a salient stimulus, and endogenous attention is the slower, voluntary, goal-directed deployment of attention. We hypothesize that exogenous attention acts primarily through the input gain and endogenous attention primarily through the feedback drive. This mapping is a testable hypothesis since the two pathways produce divergent spectral signatures (“Experimental predictions of the model”).

### Thalamo-cortical model

Alpha oscillations are a defining characteristic of thalamo-cortical dynamics, particularly prominent within the visual system during states of “relaxed wakefulness” [65, 66]. Specifically, alpha oscillations are hypothesized to be generated and modulated by thalamo-cortical loops [66, 67], suggesting that alpha rhythms signify a state of functional inhibition or “idling” in sensory pathways, potentially gated by corticothalamic feedback [67, 68]. While the hierarchical model presented in this paper was designed for cortico-cortico interactions, it may be adapted to model the thalamo-cortical interactions that contribute to the generation of alpha waves. Our theory predicts the emergence of oscillatory activity in the alpha band (8-12 Hz) under conditions of low stimulus contrast, low feedback to principal cells, and high feedback to inhibitory modulator cells (*γ* < 1, Figs. 3c,g). This aligns well with the existing literature demonstrating the role of alpha oscillations in mediating cortical inhibition [68, 69]. It is, however, unclear if the current theory will suffice, on its own, to fully capture the phenomenology of alpha waves. Furthermore, we hypothesize that the input gain modulator responses depend in part on thalamocortical loops, hence positioning the theory as a means for modeling the interplay between bottom-up sensory input and top-down feedback within the thalamocortical circuit, the gating of visual information flow, thalamic control of functional cortical connectivity [70] and thalamocortical loops in other neural systems [71].

### Limitations

Our model assumes instantaneous feedback, whereas biological feedback is delayed due to axonal conduction, synaptic transmission, and processing in higher cortical areas [72, 73]. Examining how inter-areal synchronization depends on feedback delay is an important direction for future work.

The feedforward, feedback, and recurrent weight matrices may contain both positive and negative weights. We therefore posit inhibitory interneurons to implement the sign inversion, corresponding to negative synaptic weights. These inhibitory neurons need not be one-to-one with their excitatory inputs. Rather, each may compute a partial weighted sum of their inputs corresponding to the negative terms in the matrix multiplications. Regardless, these inhibitory interneurons will introduce an additional delay. The weight matrices are thus effective (net) connectivity, and the principal cells, which are excitatory in the hypothesized mapping onto pyramidal neurons (“Biological plausibility”), interact through mixed-sign effective weights both recurrently and across areas. The circuit as written therefore does not respect Dale’s law at the level of individual connections, but it could be elaborated to do so.

Although most computations in the theory are biophysically plausible (see “Biological plausibility” and [7]), some mechanisms remain unresolved. In particular, we can only speculate about the possible cellular mechanisms for the element-wise products in Eqs. 1b and 1c (see “Biological plausibility”).

Thetime constants used in simulations are unrealistically short for individual neurons (see Table S1). Using more realistic values would shift gamma-band oscillations to lower frequencies. When all time constants were released as free parameters to fit to the experimental measurements of the power and coherence spectra (Suppl. Fig. S7), they converged to values between 0.2 and 3 ms. Multiplying all the time constants (those of the four cell types and of their firing rates, those of the noise processes, and also *α*_1,2_) by a common factor *k* would decrease the frequencies of the spectral peaks by that same factor *k*. The fixed point would be unchanged, every eigenvalue of the Jacobian would be divided by *k*, and the shape of every spectral feature would be preserved. For example, with *τ* = 10 ms the spectral peaks of Fig. 2c,e would lie at 4–6 Hz, but the rightward shift with contrast, the non-monotonic peak amplitude, the 1/ *f* ^4^ fall-off and all gain-dependent trends would be preserved. The time constants may therefore be better interpreted as effective population time constants rather than as the intrinsic time constants (capacitance/conductance) of individual neurons.

Finally, the model omits neural adaptation, a ubiquitous feature of cortical responses. Incorporating adaptation into the hierarchical framework remains an important direction for future studies.

### Biological plausibility

We hypothesize a correspondence between the variables in the theory and cortical microcircuits composed of identified cell types. Principal cells **y** are hypothesized to correspond to pyramidal neurons; inhibitory modulatory cells **a** to a combination of parvalbumin-expressing (PV+) and somatostatin-expressing (SST+) interneurons; and inhibitory interneurons **q** to vasoactive intestinal polypeptide (VIP) neurons.

In the proposed circuit (Fig. 1), Principal cells **y** excite both inhibitory modulatory cells **a** (via **u**) and inhibitory interneurons **q**. VIP-like cells **q** inhibit modulatory cells **a**, which in turn inhibit principal cells **y**. This architecture closely mirrors known cortical motifs: pyramidal neurons excite PV+, SST+, and VIP cells; VIP cells inhibit SST+ cells; and both PV+ and SST+ cells inhibit pyramidal neurons [75–79]. We further hypothesize that modulator cells **a** have large receptive fields and broad orientation-selectivity (reflecting properties of the normalization pool), consistent with the response properties of SST+ and PV+ neurons, respectively [76, 80, 81].

We also posit excitatory feedback from higher cortical areas to both principal cells **y** and modulatory cells **a** (Fig. 1). Feedback to principal cells is likely received by the distal apical dendrites of layer 1, while feedback to modulator cells **a** is commensurate with experimental evidence that feedback from higher cortical areas targets SST+ neurons [82–84], and that normalization depends in part on loops through higher visual cortical areas [57].

One potential discrepancy between the model and the cortical circuitry is that excitatory input from principal cells **y** to modulatory cells **a** is mediated by an intermediate excitatory population **u**, whereas pyramidal neurons directly excite PV+ and SST+ cells in cortex. However, the theory can be reformulated so that principal cells provide direct excitatory input to modulatory cells by substituting Eq. 1b into Eq. 1c (see Methods), without altering the model’s functional behavior.

We hypothesize that the computations performed by principal cells correspond to processes distributed across dendritic compartments of pyramidal neurons (Fig. 6). The equation governing the responses of the principal cells (see Fig. 6 and Methods, Eq. 1a) expresses the following synaptic current: (input gain) (input drive) + (recurrent gain) [recurrent drive + (feedback gain) (feedback drive)]. The input drive is computed in the distal basal dendrites, commensurate with evidence that feedforward synaptic inputs contact basal dendrites. The input gain could be controlled by inhibitory synapses near the basal trunk. The feedback drive is computed in distal apical dendrites in layer 1, which are known to receive signals from higher cortical areas. The feedback gain might be attributed to the amplification of the feedback drive within the apical trunk of pyramidal cells [85]. The recurrent drive is likely computed at more proximal apical dendrites, with recurrent gain shaped by inhibitory inputs near the proximal apical trunk.

**Figure 6:**
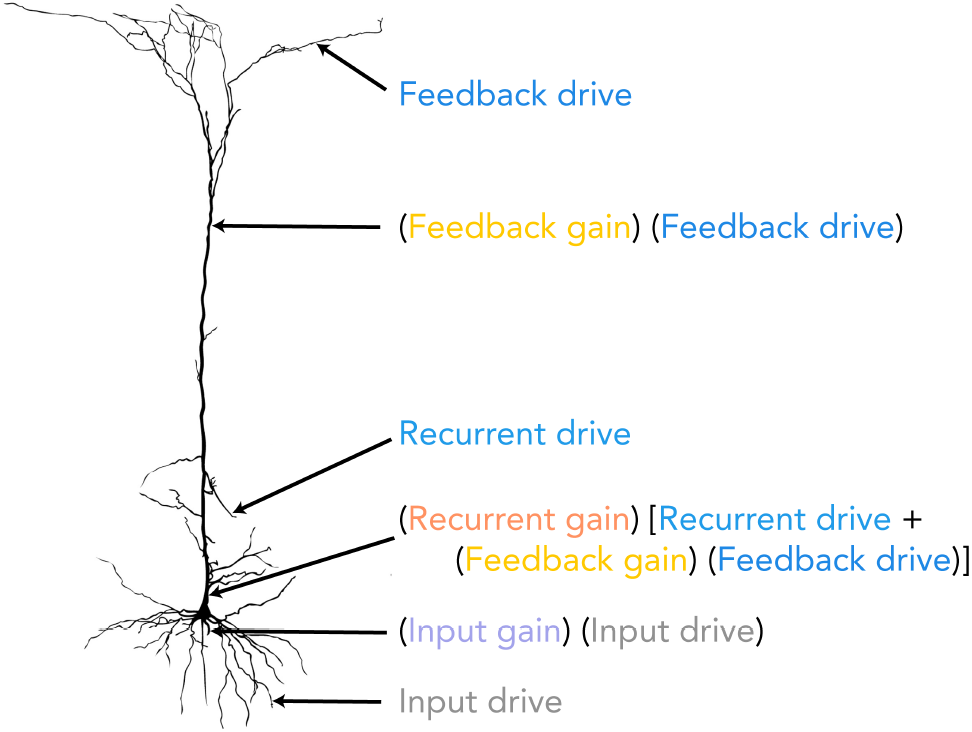
Hypothesized mapping of computational components in the model onto the dendritic compartments of a pyramidal cell. The input drive is hypothesized to be the weighted sum of feedforward inputs arriving at the distal basal dendrites. This is modulated by an input gain, corresponding to inhibition at the proximal basal dendritic trunk. In the apical tuft, the feedback drive represents a weighted sum of feedback inputs from higher cortical areas. This signal is amplified by a feedback gain within the distal apical trunk. The recurrent drive corresponds to the sum of lateral inputs on the proximal apical dendrites. The combined recurrent and feedback signals are then modulated by a recurrent gain at the proximal apical trunk. The overall synaptic current is a combination of these gain-modulated drives. The pyramidal neuron cell shown is from [74].

The weighted sums and gain control may be implemented via balanced excitatory–inhibitory push–pull and push–push mechanisms, respectively [86, 87]. Alternatively, various other mechanisms for gain control have been identified [88]. We can, however, only speculate about the possible cellular mechanisms for the element-wise products in Eqs. 1b and 1c. The element-wise product between presynaptic firing rates in Eq. 1b might be mediated by coincident spike detection, e.g., assuming that each of the spike trains is Poisson and that the synaptic current is proportional to the probability of coincident spikes within a short time window [89]. The element-wise product between postsynaptic membrane potential and presynaptic firing rate in Eq. 1c might be mediated by NMDA receptors, given that the synaptic current of an NMDA channel is jointly dependent on the presynaptic firing rate and the local membrane depolarization, in an approximately multiplicative manner [90, 91].

### Experimental predictions of the model

The theory makes several predictions that can be tested with simultaneous recordings from two or more cortical areas. For each, we outline a candidate experimental manipulation and the readouts expected. The manipulations are speculative and are offered as starting points rather than established procedures.

#### *Frequency-resolved communication subspace* (Fig. 4)

The dimensionality of the V1-V2 communication subspace is predicted to be lowest, and the prediction performance highest, in the frequency band of peak V1-V2 coherence, and this band is predicted to shift to higher frequencies with stimulus contrast. The test requires simultaneous V1 and V2 population recordings across a range of contrasts. From the cross-spectral density matrix of the recorded populations, the communication subspace, prediction performance and dimensionality can be computed at each frequency by reduced-rank regression, without band-pass filtering (Methods, “Application to experimental measurements”).

#### *Routing by differential feedback* (Fig. 5)

V1 is predicted to communicate preferentially with whichever higher area provides the stronger feedback. With simultaneous recordings from V1, V4 and V5, the relative feedback strength could be varied behaviorally, by having the subject attend to motion versus form [44, 63, 92, 93], or directly, by activating or inactivating V4 or V5 (e.g., optogenetically, electrically or pharmacologically).

#### *Normalization* (Suppl. Figs. S4 and S5)

We hypothesize that the inhibitory modulator cells **a** correspond to PV+ and SST+ interneurons (“Biological plausibility”). Manipulating these interneurons so as to weaken normalization should reduce the gamma peaks in the power and coherence spectra, remove the frequency dependence of prediction performance, and increase the dimensionality of the communication subspace.

#### *V1 input gain β*_1_ (Fig. 3, right column; Suppl. Fig. S8, middle column)

We hypothesize that the input gain is controlled by inhibitory synapses near the basal dendritic trunk of pyramidal neurons (“Biological plausibility”); identifying the interneuron type that provides this inhibition, and manipulating it, would be one way to change *β*_1_ independently of the stimulus. Increasing *β*_1_ should shift the contrast-response functions of both V1 and V2 along the log-contrast axis, shift the gamma peak to higher frequencies and narrow it, produce no alpha peak at low contrast, and decrease both inter- and within-area communication, leaving the subspace dimensionality unchanged.

#### *Feedback gain γ*_1_ (Fig. 3, left column; Suppl. Fig. S8, left column)

Feedback gain could be manipulated by activating or inactivating (e.g., by cooling or optogenetic silencing) the higher area that provides the feedback (V2 or V4), or, more speculatively, by targeting the amplification of the feedback drive in the apical trunk of pyramidal neurons (“Biological plausibility”). Increasing *γ*_1_ should produce a larger change in contrast gain in V2 than in V1; at low contrast, the alpha peak in V1 power and the beta peak in V1-V2 coherence that are present at low feedback gain should vanish; the gamma peak should shift to higher frequencies and broaden; and inter-areal communication should increase while within-area communication decreases, with the subspace dimensionality unchanged.

#### *V2 input gain β*_2_ (Suppl. Fig. S8, right column)

The V2 input gain scales the V1 → V2 drive. It could be manipulated by targeting the V1 → V2 projection (e.g., optogenetic modulation of V1 axon terminals in V2) or the corresponding basal-trunk inhibition of V2 pyramidal neurons, with V1 and V2 recorded simultaneously. Its signature differs from those of the other two gains: V1 responses are unchanged while V2 responses are enhanced; the gamma peak shifts to higher frequencies, broadens, and decreases in amplitude; there are no alpha or beta peaks at low contrast; and inter-areal communication increases while within-area communication is unchanged.

## Methods

### The Model

We propose a multistage hierarchical recurrent circuit that incorporates feedforward connections from the primary visual cortex (V1) to higher visual areas (e.g., V2, V4, MT) and feedback connections from these higher brain areas to V1 (Fig. 1). We refer to V2 throughout this paper for concreteness, but it is merely an example of a higher brain area that is reciprocally connected with V1. This model builds on the ORGaNICs framework developed by Heeger et al. [6, 7].

The response of principal neurons in the model is a sum of three key components: i) input drive, a weighted sum of neural responses from a brain area that provides feedforward input (e.g., LGN → V1 and V1 → V2), modulated by input gain; ii) feedback drive, a weighted sum of neural responses from the higher cortical area (e.g., V1 ← V2 or V1 ← V4), modulated (viz., scaled) by feedback gain; and iii) the sum of recurrent drive, a weighted sum of neuronal responses from other neurons in the same brain area (e.g., V1 → V1 and V2 → V2) and scaled feedback drive, collectively modulated by recurrent gain.

The proposed neural circuit comprises four distinct cell types: the principal neurons (**y**), the modulatory excitatory neurons (**u**), the modulatory inhibitory neurons (**a** and **q**), with the following dynamics:

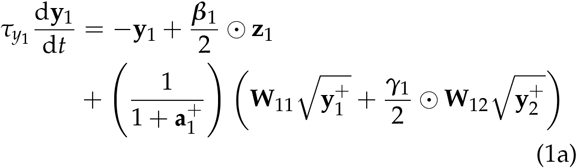

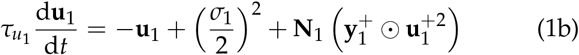

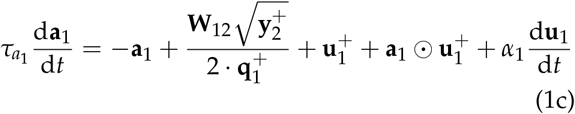

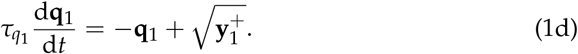

The above equations are for V1; V2 follows a similar set of equations (see *SI Appendix*). The membrane potentials of neurons in cortical areas V1 and V2 are denoted by **y**_1_ and **y**_2_, respectively, while their corresponding firing rates are represented by 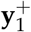 and 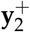. The ⊙ symbol represents the Hadamard product, element-wise multiplication of vectors. To estimate the firing rates 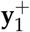 and 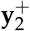 from their respective membrane potentials, we employ a modified Gaussian Rectification (GR) model [94, 95]. The variable **z**_1_ denotes the input drive to V1 from the preceding brain area (the lateral geniculate nucleus of the thalamus, LGN), computed as a weighted (**W**_*zx*_) sum of the LGN firing rates (Fig. 7a illustrates the resulting orientation tuning curves). *τ*_**k**_ is the time constant for each subtype of neuron, where 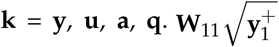 and 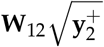 are the recurrent and feedback drive, respectively, where **W**_11_ is the recurrent weight matrix for V1. **W**_11_ has a center-surround structure where the closer connections are excitatory and the more distant connections are inhibitory (Fig. 7b), same as in [7]. The recurrent weight matrix for V2, **W**_22_, has a similar structure. The maximum eigenvalue for these recurrent weight matrices is set to unity to ensure stability. However, we have shown that this need not be the case because normalization ensures stability [8, 9]. **W**_12_ represents the inter-areal feedback connectivity matrix from V2 to V1; it is diagonally dominant, with net excitatory entries near the diagonal and weak net inhibitory entries farther from it (Fig. 7c). The feedforward matrix from V1 to V2, **W**_21_, is taken to be the transpose of **W**_12_. Given that **W**_12_ is symmetric, it follows that **W**_12_ = **W**_21_. The normalization matrix in V1 (**N**_1_) and V2 (**N**_2_) is taken to be all ones, meaning each neuron contributes equally to the normalization. All of these matrices describe effective connectivity; negative weights may, for example, be implemented with disynaptic inhibition through interneurons that are not modeled explicitly (see Discussion). The input gain and feedback gain for V1 are modulated by *β*_1_ and *γ*_1_, respectively. The recurrent and feedback amplifications in V1 are modulated dynamically by the inhibitory neuron population **a**_1_.

**Figure 7:**
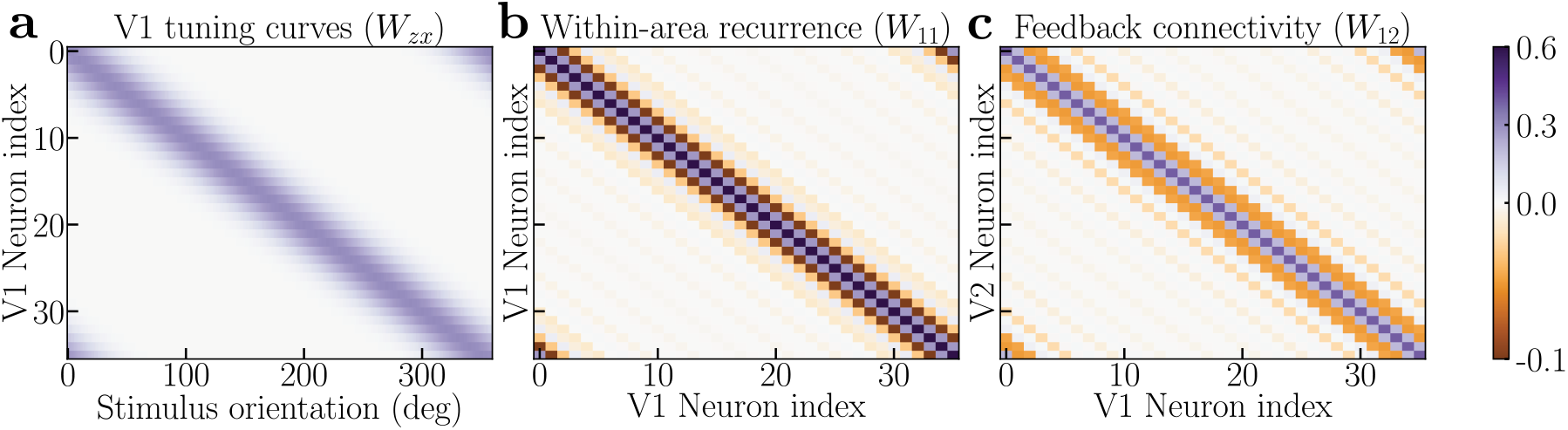
Connectivity matrices in the hierarchical recurrent circuit model. **a**, Orientation tuning curves corresponding to the V1 encoding matrix (**W**_zx_). Each row represents the tuning curve of a V1 neuron. The weight matrix itself has positive and negative synaptic weights like standard models of orientation-selective receptive fields. **b**, The recurrent connectivity matrix for V1 (**W**_11_). The V2 recurrent connectivity matrix **W**_22_ has a similar structure. **c**, The Feedback connectivity matrix from V2 to V1 (**W**_12_). The corresponding feedforward matrix from V1 to V2, (**W**_21_), is its transpose. Since **W**_21_ is symmetric, the feedforward and feedback matrices are identical (**W**_12_ = **W**_21_).

Both the V1 and V2 neurons follow the normalization equation *exactly* when *β* and *γ* are set to unity (i.e. when there is identical feedback to principal neurons (**y**_1_) and inhibitory modulator (**a**_1_) neurons) and **W**_11_ is the identity matrix (i.e., each neuron recurrently amplifies itself), but the circuit closely approximates normalization for a wide range of parameters. For V1 neurons, normalization is expressed as:

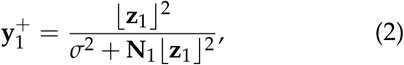

and likewise for V2

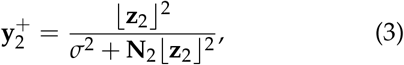

where 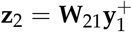 is the drive from V1 to V2, *σ* is the semi-saturation constant. Normalization is achieved dynamically via recurrent amplification (amplifying weak inputs but not strong inputs) with the three modulatory neurons, **u**_1_, **a**_1_ and **q**_1_ whose dynamics are governed by Eq. 1b, Eq. 1c, and Eq. 1d respectively. The equations for modulatory neurons are derived by substituting the normalization equation (Eq. 2) in the equation for the principal neurons (Eq. 1a) (see *SI Appendix* for derivation). An increase in the activity of principal neurons (**y**_1_) leads to enhanced activation in **u**_1_ and **q**_1_. This, in turn, drives a net increase of activity in modulatory neurons (**a**_1_). The elevated **a**_1_ activity subsequently exerts an inhibitory effect on **y**_1_, thereby implementing normalization by the recurrent interactions between the neural populations (see Appendix for derivation).

The parameter *γ*_1_ modulates the feedback gain to the principal neuron population (**y**_1_). This determines the relative feedback to the principal neurons (**y**_1_) compared to the inhibitory modulator neurons (**a**_1_). As a rule of thumb, *γ*_1_ < 1 indicates greater feedback inhibition, *γ*_1_ = 1 denotes a balance between feedback excitation and inhibition, and *γ*_1_ > 1 signifies more feedback excitation. For simplicity, we have set the feedback to the inhibitory neurons (**a**_1_) at a fixed value of 0.50 (hence, the factor of 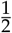 in equation Eq. 1c and Eq. 1a, see *SI Appendix*). Similarly, the parameter *β*_1_ modulates the relative input gain to the principal neuron population (**y**_1_) compared to the modulatory excitatory neuron population (**u**_1_). For simplicity, we set the input gain to the modulatory neurons (**u**_1_) to 0.50 (hence, the factor of 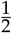 in equation Eq. 1a and Eq. 1b). Note that when *β*_1_ and *γ*_1_ are both set to unity and the recurrent matrix is identity, the network achieves excitatory-inhibitory balance exactly, and the normalization (Eq. 2) is followed precisely.

We modeled each brain area using a ring model, where principal neurons are arranged in a circle to represent continuous features like stimulus orientation. The recurrent connectivity between these neurons is determined by the distance between them (Fig. 7b). Each modeled area consists of 72 neurons for each of the four specified types (**y**_1_,**u**_1_,**a**_1_,**q**_1_). To simulate sensory input, a visual stimulus with a defined contrast and orientation is presented. The input drive is the product of the stimulus vector with the encoding matrix, whose rows are the tuning curves of the V1 principal neurons (Fig. 7a). The tuning curves are modeled as raised cosine functions, each peaking at the neuron’s preferred orientation.

### Numerical computation of fixed points

For all results reported here, the fixed point was computed numerically by integrating the deterministic dynamics to steady state. Starting from small uniform initial conditions, the dynamics were integrated with an adaptive Runge–Kutta scheme (RK45; relative and absolute tolerances of 10^−10^) over a window spanning several thousand times the slowest time constant. The steady state was accepted when every state variable changed by less than 10^−6^ between consecutive output samples, and the Jacobian at the fixed point was verified to have eigenvalues with negative real parts (i.e., a stable fixed point).

The numerically computed fixed points closely approximate Eq. 2, to within 0.6% with the center-surround recurrent weight matrix (not the identity) and the feedback gain *γ*_1_ set to one. The deviation from the normalization fixed point increases as the feedback gain *γ*_1_ and the input gain *β*_1_ deviate from 1.0.

### Power and coherence spectra

The time evolution of neural population activity in V1 and V2 can be described by a general stochastic differential equation (SDE) with an additive noise term **L**d**W**:

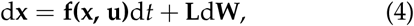

where **x** ∈ ℝ^*n*^ is a vector representing the activity of neurons; **f(x, u)** ∈ ℝ^*n*^ represents the deterministic part of the dynamics explained above (Eq. 1); d**W** is a vector of independent Gaussian increments with correlation matrix **D** (E[d**W** · d**W**^T^]), and **L** ∈ ℝ^*n*×*n*^ is the dispersion matrix defining how noise enters the system. Given the fixed point of the dynamical system, we can linearize the system around the fixed point (**x**^∗^):

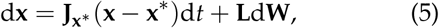

where **J**_**x**_∗ is the Jacobian computed at the fixed point **x**^∗^. This linearization leads to a wide-sense stationary Gaussian process, **x**(*t*), whose power spectral density matrix, ***S***(*ω*) ∈ ℝ^*n*×*n*^, is given by [96]:

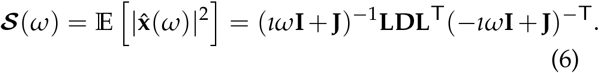

From ***S***(*ω*), we can directly compute the coherence, *κ*_*ij*_, between any two *i* and *j* neurons in the two areas:

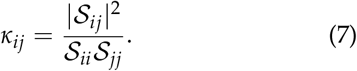

The linearization was validated by direct numerical integration of the full nonlinear stochastic system, including all noise sources, with the Euler–Maruyama scheme (time step 10 *µ*s; 100 independent trials of 16 s after a 1 s burn-in, at each contrast); power and cross-spectra were estimated from the simulated trajectories by Welch’s method (4 s segments, 50% overlap) and compared with Eqs. 6 and 7 (Suppl. Fig. S6; *SI Appendix*, “Numerical validation of the analytical spectra”). A quantitative comparison with experimental spectra, in which the model parameters are fitted, is described in the *SI Appendix* (“Quantitative fit to experimental power and coherence spectra”) and shown in Suppl. Fig. S7.

### Communication subspaces

Here we present the method used to characterize the communication subspaces between two subpopulations (source and target) of neurons [15]. These subpopulations may comprise distinct neurons in the same brain area or across different brain areas. The basic idea is to determine how well the fluctuations in responses of a target subpopulation can be predicted using a linear transform (a weighted sum) of the fluctuations in responses of the source subpopulation, and to determine the dimensionality of this linear transform.

The fluctuations in the responses and their co-variability are fully determined by the correlation matrix at steady-state when the strength of the noise is small. Therefore the correlation matrix, denoted as **C**(0), for the responses **x**(*t*) can be computed by solving the Lyapunov equation [96]:

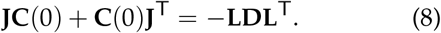

Once **C**(0) is obtained, we follow the method proposed by Semedo et al. [15] to construct an analytical model of the communication subspace. To achieve this, we divide the mean-subtracted responses of the entire neuronal population, denoted as **x**, into two subsets: **s** (responses of the source neurons, e.g., in V1) and **t** (responses of the target neurons, e.g., in V2 or V1). We consider a linear model for predicting target responses using the source responses, 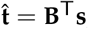, where the mean squared error (MSE) is given by:

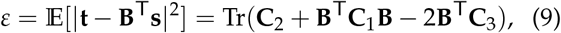

where **C**_1_ = E[**ss**^T^], **C**_2_ = E[**tt**^T^], and **C**_3_ = E[**st**^T^]. These matrices can be directly obtained from **C**(0). To find the optimal readout matrix **B**_opt_, we minimize *ε* with respect to **B**, leading to the Ordinary Least Squares solution:

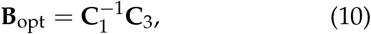

First, we define the covariance of the target activity *predicted* by the optimal model (note the “hats”), which we denote as **Ĉ**_2_:

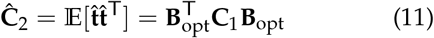

By substituting the expression for **B**_opt_, we arrive at a key relationship:

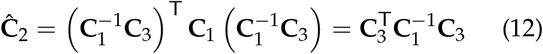

The principal dimensions of this predicted covariance matrix, **Ĉ**_2_, define the communication subspace. These are found by computing the eigen-decomposition:

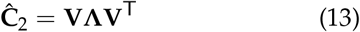

Here, the columns of the orthogonal matrix **V** are the eigenvectors of **Ĉ**_2_ and **Λ** is a diagonal matrix of the corresponding eigenvalues, sorted in descending order. The prediction performance for a reduced-rank approximation of dimension *i* is defined as: A rank-*i* approximation, **B**_RRR_, is constructed by projecting the optimal solution **B**_opt_ onto the subspace spanned by the top *i* eigenvectors of **Ĉ**_2_.

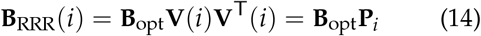

where **V**(*i*) is a matrix containing first *i* columns of **V**. Here **P**_*i*_ = **V**(*i*)**V**^T^(*i*) is the projection matrix. Note that **P**_*i*_ is symmetric 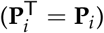 and idempotent 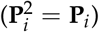. We define the prediction performance as:

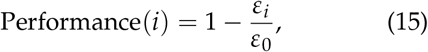

where *ε*_*i*_ is the MSE for the rank-*i* approximation

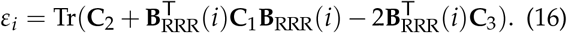

*ε*_0_ is the MSE for the case where the readout matrix is a null matrix, corresponding to the total variance of the target population (*ε*_0_ = Tr(**C**_2_)).

To simplify the expressions for *ε*_*i*_. We substitute **B**_RRR_(*i*) = **B**_opt_**P**_*i*_ into the expression:

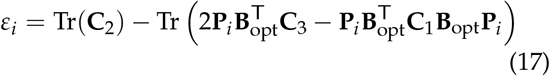

Now, we substitute 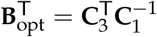 and recognize that the term 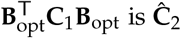 is **Ĉ**_2_. Using the cyclic property of the trace (Tr(**AB**) = Tr(**BA**)) and the idempotent property of the projection matrix 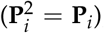, we can simplify further:

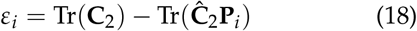

By applying the cyclic property and expanding **P**_*i*_, we can see that the second term is the sum of the top *i* eigenvalues (*λ*_*j*_) of **Ĉ**_2_.

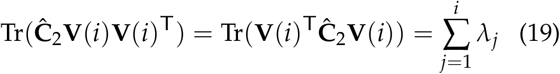

Now the expression for prediction performance simplifies to:

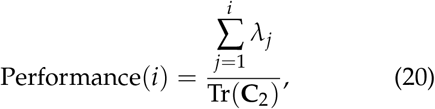

Which can also be interpreted as:

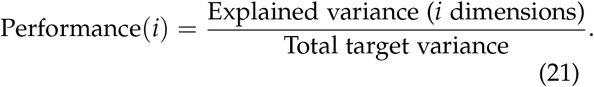

The maximum prediction performance (full-rank approximation) is given by:

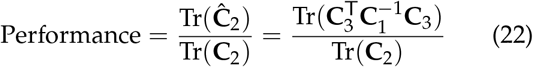

For the communication-subspace analyses (Figs. 2g and 3k,l), source and target populations of 30 principal neurons were drawn at random from the 72 neurons of each area, and prediction performance and dimensionality were averaged over random draws. The dependence of the dimensionality on the number of modeled and sampled neurons is characterized in Suppl. Fig. S9.

#### Interpretation of the results

The prediction performance (Eq. 22) is governed by the interplay between the source (**C**_1_), target (**C**_2_), and cross-population (**C**_3_) covariances. The effectiveness of communication (indicated by a large predictive performance) can be understood through three key factors: 1) *Source-target correlation*: Prediction performance is directly enhanced by a stronger correlation between the source and target populations (i.e., larger **C**_3_ norm). 2) *Signal reliability*: Prediction performance is boosted by the reliability of the signals. Less noisy activity in source and target populations, indicated by smaller diagonal entries in **C**_1_ and **C**_2_, leads to better prediction performance. 3) *Alignment*: Prediction performance is maximized when the principal directions of source covariance (eigenvectors of **C**_1_) align with the principal directions of source-target covariance (left singular vectors of **C**_3_).

This framework also explains the results in the frequency domain. The key metric here is the cross-power spectrum, which is the Fourier transform of the cross-covariances, and coherence, which is the squared cross-power spectrum normalized by the product of the auto-spectra, Eq. 7. At frequencies of high coherence, i.e., of large cross-covariance between source and target relative to their auto-covariances, the prediction performance is necessarily highest as well, since both quantities are computed from the same cross-spectral density matrix.

The dimensionality of the communication subspace is given by the rank of the matrix 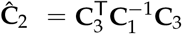. Since the source covariance matrix **C**_1_ is a full-rank matrix, the dimensionality of the subspace is governed by the rank of the cross-covariance matrix, **C**_3_. In the frequency domain, this means the dimensionality at a specific frequency is determined by the rank of the cross-power spectral density matrix (see Suppl. Fig. S5a). Our results show that the extent of normalization from surrounding neurons plays a critical role in shaping the dimensionality of the communication subspace. With surround normalization, dimensionality decreases specifically at frequencies where coherence is strongest (Fig. 4c). Normalization reduces dimensionality because when just a small subset of neurons is strongly driven, and their responses are highly correlated across areas (V1 and V2), normalization suppresses the activity of surrounding neurons with weaker drive. This suppression, resembling a soft winner-take-all process, diminishes the contribution of those less active neurons to the population’s shared variability, leading to a lower effective dimensionality.

In contrast, without this normalization, no such correlation between coherence and dimensionality is observed (Fig. S4h).

### Frequency decomposition of the communication subspace

To analyze how communication between different brain areas varies across frequencies, we consider the frequency decomposition of the neural responses. The Wiener-Khinchin theorem states that the autocorrelation function **C**(*τ*) is the inverse Fourier transform of the power spectral density (PSD) matrix (***S***(*ω*)):

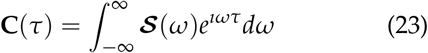

To isolate **C**(*τ*) at a particular frequency (*ω*^∗^), we apply an idealized narrow band-pass filter centered at *ω*^∗^. Specifically, the filter’s transfer function *H*(*ω*) is given by,

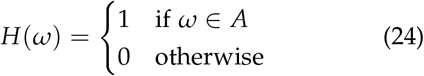

where *A* = [*ω*^∗^ − *ϵ, ω*^∗^ + *ϵ*] ∪ [ − *ω*^∗^ − *ϵ*, − *ω*^∗^ + *ϵ*] and *ϵ* is a small positive value defining the narrow bandwidth 2*ϵ* around *ω*^∗^. The spectral density matrix of this filtered signal is given by,

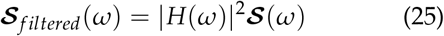

The resulting autocorrelation of the filtered signal is expressed as:

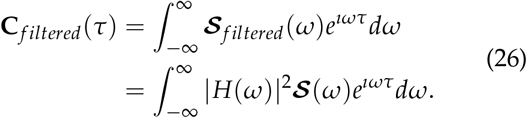

Using the definition of *H*(*ω*) and assuming *ϵ* to be sufficiently small such that (*ω*) and *e*^*ıωτ*^ are approximately constant over the integration intervals of width 2*ϵ*, the frequency-specific autocorrelation at *ω*^∗^ is:

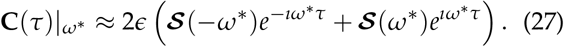

At zero time lag (*τ* = 0),

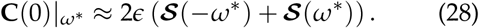

Since the PSD is Hermitian, i.e., ***S*** (*ω*) = ^∗^ ***S*** (− *ω*)), we have

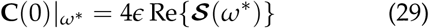

This frequency-specific autocorrelation matrix (**C**(0)|_*ω*_∗) can then be used in place of **C**(0) in the communication subspace analysis described above to evaluate the prediction performance at any frequency. As our analysis does not depend on the absolute scaling of this matrix, the prefactor 4*ϵ* can be disregarded, and we effectively use Re {***S*** (*ω*^∗^)} as the frequency-specific correlation matrix.

#### Application to experimental measurements

The same construction applies directly to experimental recordings, and no band-pass filtering or separate regression per frequency band is required. Given simultaneously recorded source (e.g., V1) and target (e.g., V2) population activity (spike counts or LFP), one estimates the cross-spectral density matrix ***S***(*ω*) of the pooled population (e.g., by Welch’s or the multitaper method). At each frequency *ω*^∗^, Re {***S*** (*ω*^∗^)} is the frequency-specific covariance matrix; partitioning it into the source, target and cross blocks **C**_1_, **C**_2_ and **C**_3_ gives 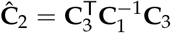, the prediction performance at each rank (Eq. 20), and the dimensionality, i.e., the number of eigenvalues of **Ĉ**_2_ required to reach the full-rank performance (Eq. 22) within a fixed tolerance. Repeating this at every frequency yields the experimental counterparts of Fig. 4b,c. The prediction to be tested is that the dimensionality is smallest at the frequency of peak V1-V2 coherence and that this frequency increases with contrast.

## Code Availability

The code to generate all the figures can be found in the following GitHub repository: https://github.com/asit-pal/Hierarchical-ORGaNICs

## Acknowledgements

We thank Pascal Fries and Adam Kohn for generously providing the experimental data in Fig 2. We acknowledge Michael Halassa, Hillel Adesnik, Adam Kohn and Flaviano Morone for their valuable feedback on the manuscript, and Ruben Coen-Cagli, John Maunsell, and Jason MacLean for insightful discussions. This work was initiated in part at the Aspen Center for Physics, which is supported by National Science Foundation grant PHY-2210452. This work was supported by the National Institute of Health under award numbers R01MH137669 and R01EY035242. S.M. also acknowledges support from the Simons Center for Computational Physical Chemistry (Simons Foundation grant 839534, MT). This work was supported in part through the NYU IT High-Performance Computing resources, services, and staff expertise.

