## Supplementary Information for "Hierarchical Neural Circuit Theory of Normalization and Inter-areal Communication"

**Contents**

I. Additional theoretical predictions 1

II. Numerical validation of the analytical spectra 4

III. Quantitative fit to experimental power and coherence spectra 5

IV. Robustness with respect to model parameters 6

V. Derivation of the General Hierarchical Recurrent Normalization Circuit 9

VI. Noise Model 13
A. Intrinsic neural circuit noise 13
B. Extrinsic noise in the local field potential 14

References 14

### I. Additional theoretical predictions

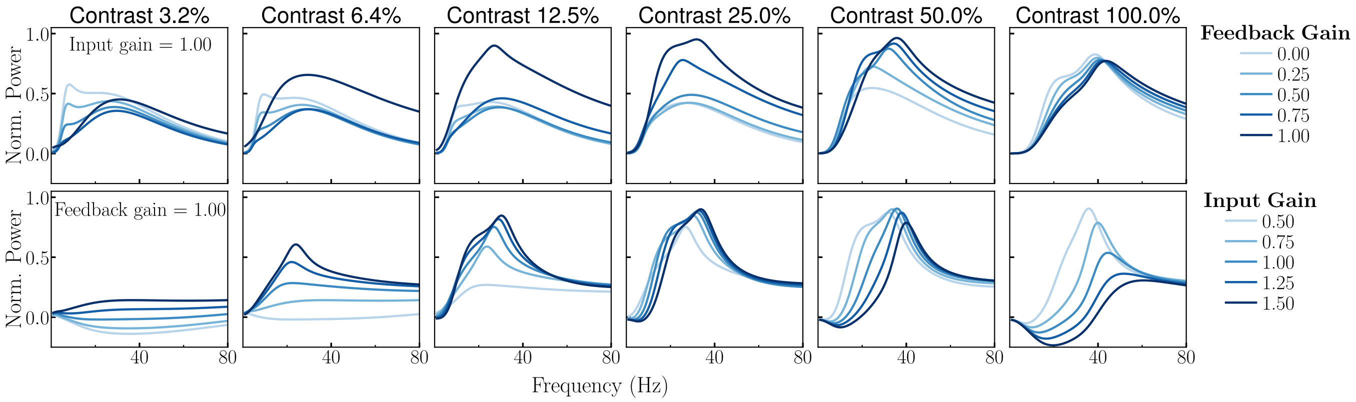


FIG. S1. **Theoretical predictions of varying feedback and input gain on power spectra across different contrast levels.** Each column displays results for a specific contrast, with contrast increasing from left to right across panels. **Top row,** Normalized power as a function of frequency, for varying feedback levels (indicated by the blue line color intensity), with input gain kept at baseline (1.0). **Bottom row,** Normalized power as a function of frequency for varying input gain levels, with feedback gain kept at baseline (1.0). In all panels the V2 input gain $\beta_2$ is held at its baseline value (1.0).

*
†
‡

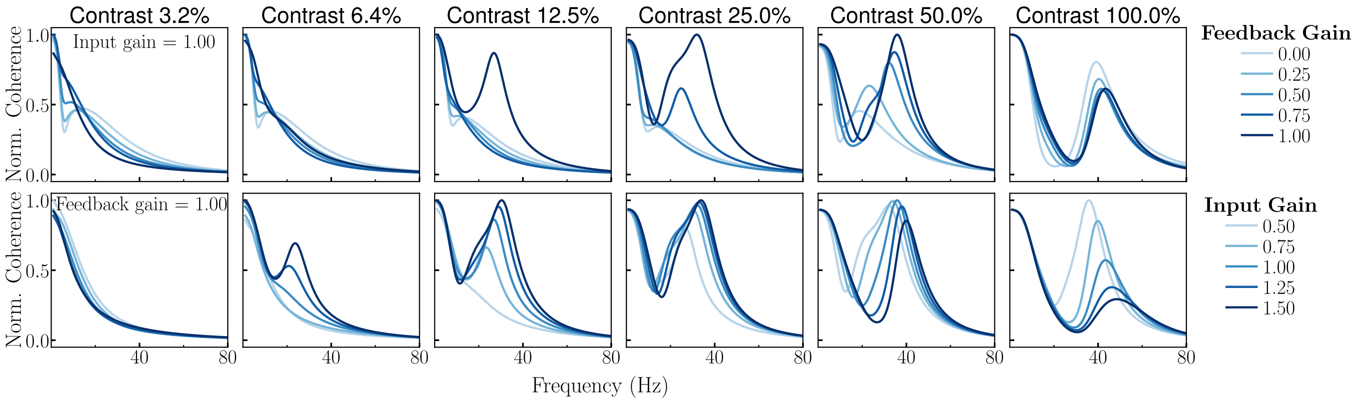


FIG. S2. **Theoretical predictions of varying feedback and input gain on V1-V2 coherence across different contrast levels.** Each column displays results for a specific contrast, with contrast increasing from left to right across panels. **Top row,** Normalized coherence as a function of frequency, for varying feedback levels (indicated by the blue line color intensity), with input gain kept at baseline (1.0). **Bottom row,** Normalized coherence as a function of frequency for varying input gain levels, with feedback gain kept at baseline (1.0). In all panels the V2 input gain $\beta_2$ is held at its baseline value (1.0).

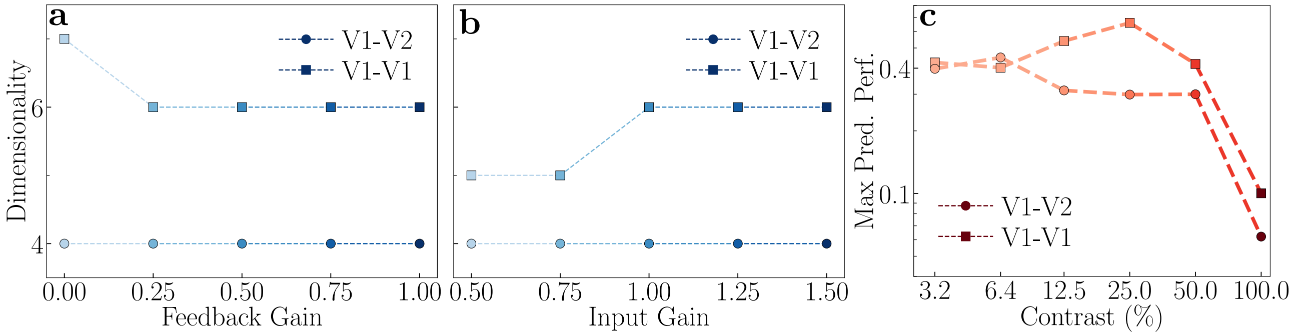


FIG. S3. **Theoretical predictions: Varying feedback gain, V1 input gain, and contrast on communication subspaces. a,** Dimensionality of inter-areal (V1-V2, circles) and within-area (V1-V1, squares) communication subspaces as a function of feedback gain, at 100% stimulus contrast. **b,** Dimensionality of inter-areal (V1-V2, circles) and within-area (V1-V1, squares) communication subspaces as a function of V1 input gain, at 100% stimulus contrast. **c,** Maximum prediction performance for inter-areal (V1-V2, circles) and within-area (V1-V1, squares) communication as a function of stimulus contrast. In all panels the V2 input gain $\beta_2$ is held at its baseline value (1.0).

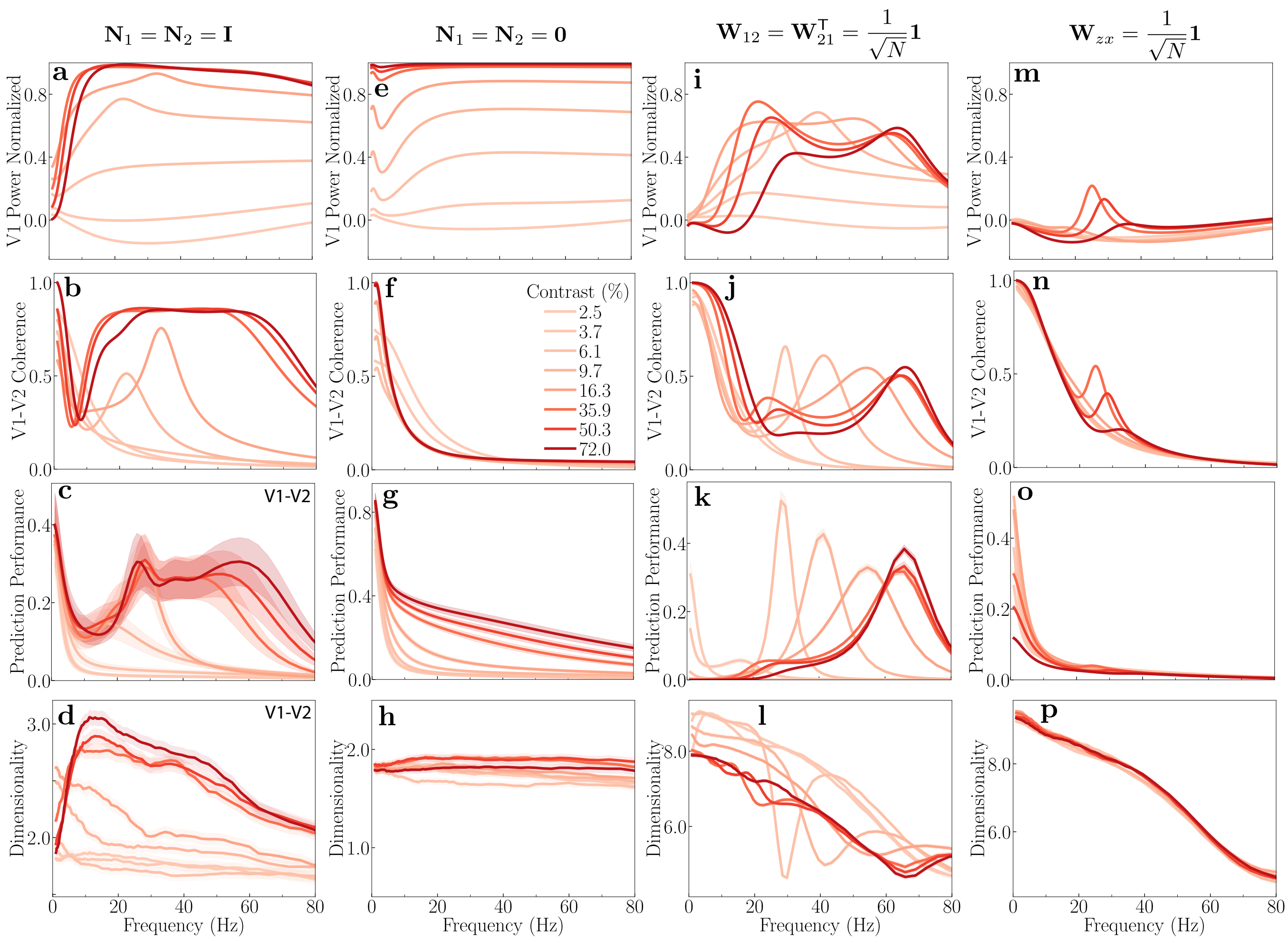


FIG. S4. **Effect of normalization and V1-V2 connectivity on oscillatory activity and communication subspaces.** The impact of divisive normalization and inter-area connectivity on network dynamics is explored across four model configurations (columns). For each model, we plot the V1 power spectrum, V1-V2 coherence, prediction performance, and the dimensionality of the communication subspace (rows). For communication subspace analysis, 18 out of 72 simulated neurons were randomly selected from each area, and the analysis was repeated multiple times; shaded areas represent the standard error of the mean (SEM). The number of neurons for the communication subspace analysis was selected for direct comparison with published experimental measurements of communication subspaces with broadband (natural image) input stimuli (see Suppl. Fig. 7 of [1]). **a-d, Self-normalization**: Model predictions when the normalization matrix is set to identity ($\mathbf{I}$), meaning each neuron only normalizes its own activity. **a,** Normalized V1 power spectra at different stimulus contrast values. **b,** The coherence peaks between V1 and V2 are narrow band at low contrast, and become broadband at high contrasts ($> 16\%$). **c,** Prediction performance is highest at frequencies corresponding to peak coherence. **d,** The communication subspace dimensionality is low, but increases with contrast. **e-h, No-normalization**: Model predictions when the normalization matrix is a null matrix (i.e, normalization is removed). **e,f** Removing normalization eliminates prominent oscillatory activity in both the power and coherence spectra. **g,** Removing normalization also eliminates peaks in prediction performance. **h,** Subspace dimensionality remains low and shows little or no dependence on frequency or contrast. **i-l, Uniform inter-area connectivity**: Model prediction when the V2 to V1 connectivity matrix modified to $\mathbf{I} + \frac{1}{\sqrt{N}}\mathbf{1}$, where N is number of principal neurons and $\mathbf{1}$ is an ($N \times N$) matrix of ones. **i,** V1 power shows distinct peaks in the low and high gamma frequency range. **j-k,** High coherence corresponds to high prediction performance. **l,** The dimensionality is higher compared to when inter-area connectivity is sparse, but still much lower than the number of neurons (18), and still markedly reduced at frequencies where coherence is strongest. **m-p, Uniform external input**: V1 tuning curves are modified to be flat, signifying an equal preference for all stimulus orientations (or equivalently, that the input stimulus is broadband, e.g., noise or natural images). **m,n,** Oscillatory activity is evident in both power and coherence spectrum. **o,** Prediction performance decreases with frequency. **p,** Communication subspace dimensionality is higher compared to when the tuning curves (or stimulus) is narrow band, but still much lower than the number of neurons (18).

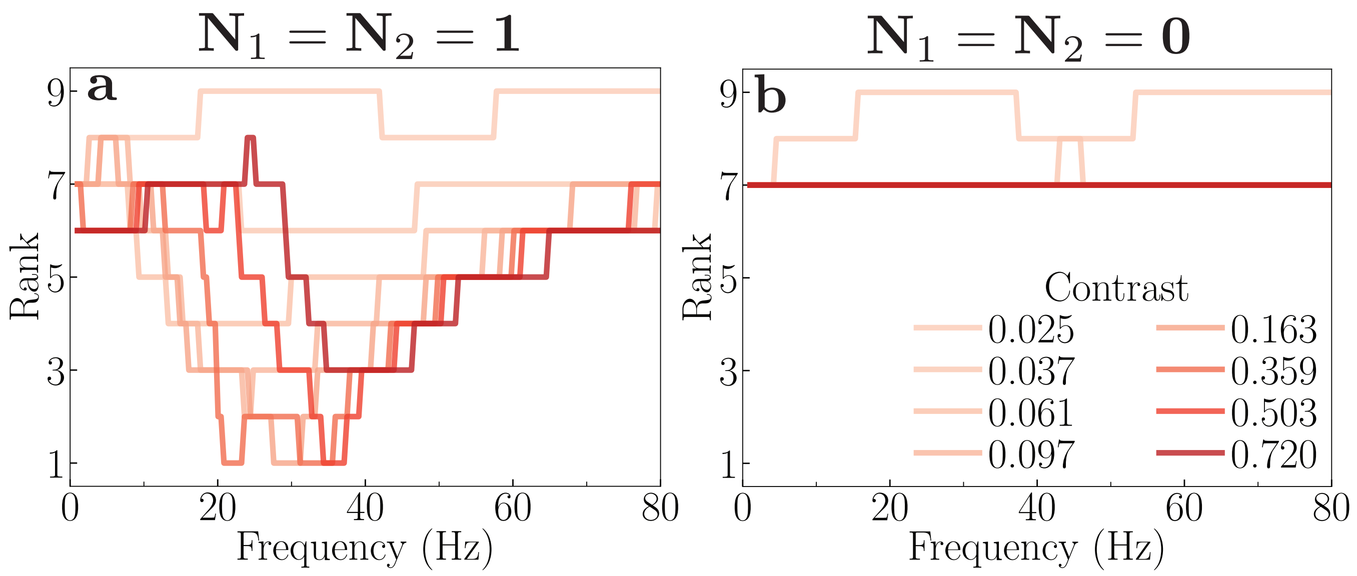


FIG. S5. **Effect of normalization on the rank of the V1–V2 cross-covariance matrix across frequencies. a,** Normalization matrices for V1 and V2 are set to be all ones, implying that each neuron in the same area contributes equally to the normalization pool. The rank follows trends similar to the dimensionality (see Fig. 4c); the rank decreases at the coherent frequencies. **b,** Normalization matrices for V1 and V2 are set to zeros, corresponding to no normalization. The rank of the cross-covariance matrix shows no frequency-specific behavior.

### II. Numerical validation of the analytical spectra

The power and coherence spectra reported in the main text are obtained by linearizing the stochastic dynamics about the fixed point (main text, Methods, "Power and coherence spectra"). To verify this linearization, we integrated the full nonlinear stochastic system, i.e., the membrane-potential dynamics (Eqs. S31–S38), the firing-rate dynamics with firing-rate noise (Eq. S43), and the filtered synaptic noise (Eq. S44), with the Euler–Maruyama scheme at the baseline parameters (Table S1) and noise parameters (Table S3). Trajectories were initialized at the deterministic fixed point and integrated with a time step of $10\,\mu$s (the fastest mode of the linearized system has a time constant of about 0.5 ms); after a burn-in of 1 s, 100 independent trials of 16 s were recorded at each contrast of the sweep shown in Fig. 2c,e (0%, 2.5%, 3.7%, 6.1%, 9.7%, 16.3%, 35.9%, 50.3%, and 72%). Power spectra and cross-spectra of the membrane potentials were estimated by Welch's method (segments of 4 s, 50% overlap) and averaged over trials; the extrinsic LFP noise of Sec. VI B, which is independent of the circuit dynamics, was added analytically to the estimated spectra, exactly as in the analytical calculation. The simulated power spectra were normalized by the simulated spectrum at 0% contrast, with the same normalization as in the main text, and the coherence was computed between the maximally driven principal neurons in V1 and V2 (main text, Eq. 7). The comparison with the closed-form expressions is shown in Fig. S6: across all contrasts, the root-mean-square deviation is below 0.017 for the normalized power and below 0.018 for the coherence, and the maximum absolute deviation is below 0.072, confirming that the linearization is accurate at the baseline parameters. The linearization holds for small fluctuations about the fixed point, and the simulations bear this out. When the amplitudes of the noise sources are increased beyond the values of Table S3, the deviation between the simulated and the analytical spectra grows, and it does so first at the lowest contrasts ($< 10\%$): there the activities lie near zero, the noise fluctuations can drive them negative, and the rectification clips them to zero, which the linearized dynamics do not capture. Changing the absolute noise amplitude does not change the analytical results, since the plotted spectra are normalized. The deviations do not, however, grow with the gains: changing the input or feedback gain changes the fixed point, and the linearization about the new fixed point remains valid, because its accuracy is controlled by the strength of the fluctuations relative to the fixed point rather than by the gains themselves. All results reported in the paper use the baseline parameters, for which the linearization is accurate at every contrast and for all reported gain values; the stable ranges of the gains are given in Table S2.

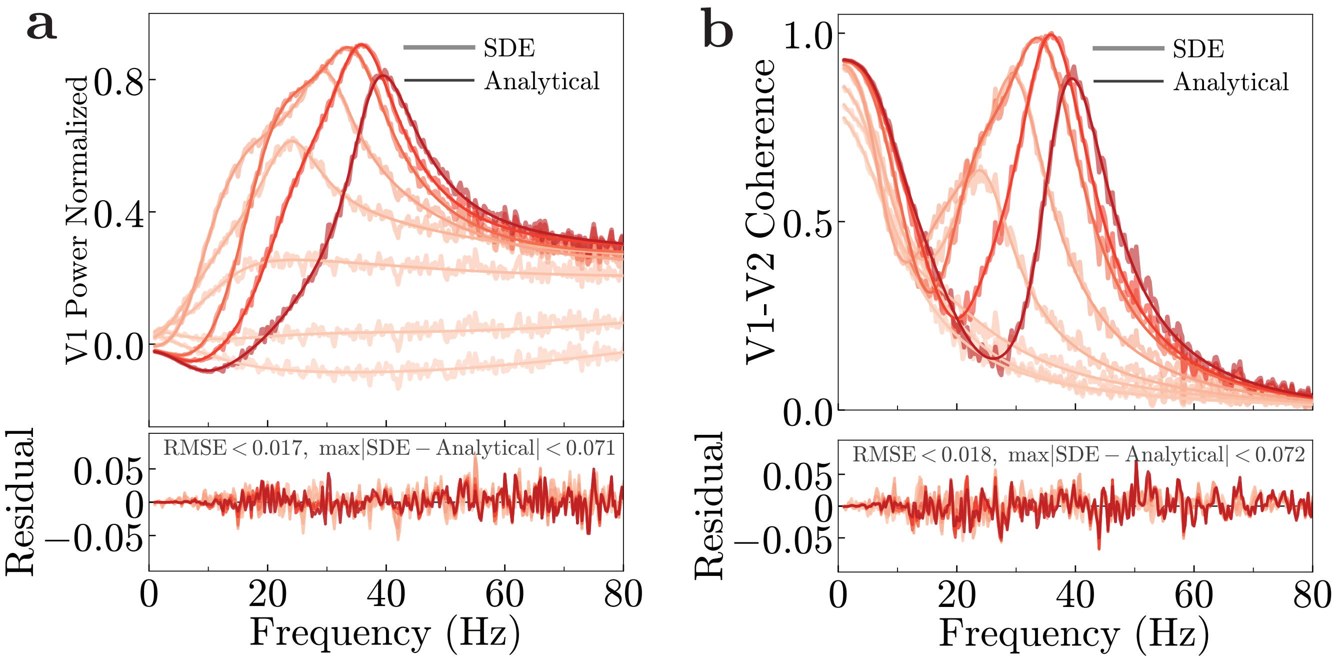


FIG. S6. **Validation of the analytical power and coherence spectra against direct stochastic simulation.** The full nonlinear stochastic system (main text, Methods, "Power and coherence spectra"; noise model of Secs. VI A and VI B) was integrated numerically, and the spectra were estimated directly from the simulated trajectories (light, jagged lines; "SDE") and compared with the closed-form expressions obtained by linearizing the dynamics about the fixed point (dark, smooth lines; "Analytical"). Line color indicates stimulus contrast (increasing from light to dark); all other parameters are the baseline values (Table S1). **a,** Normalized V1 power spectra. **b,** V1–V2 coherence spectra. Bottom panels show the residual (SDE − Analytical) at each contrast. Across all contrasts, the root-mean-square error is below 0.02 and the maximum absolute deviation is below 0.08 (power: RMSE $< 0.017$, $\max|\text{SDE} - \text{Analytical}| < 0.071$; coherence: RMSE $< 0.018$, $\max|\text{SDE} - \text{Analytical}| < 0.072$), confirming that the linearization is accurate in this parameter regime.

### III. Quantitative fit to experimental power and coherence spectra

All comparisons between theory and experiment in the main text use a single, unfitted set of baseline parameters. To quantify how well the theory can account for experimentally measured spectra, we fit the model to the V1 power spectra and V1–V2 coherence spectra of Roberts et al. [2] (digitized from the published figures at eight contrast levels: 2.5%, 3.7%, 6.1%, 9.7%, 16.3%, 35.9%, 50.3%, and 72%). The fitted model is the same circuit as in the main text, with the connectivity matrices ($\mathbf{W}_{zx}$, $\mathbf{W}_{11}$, $\mathbf{W}_{22}$, $\mathbf{W}_{12} = \mathbf{W}_{21}^{\intercal}$), the all-ones normalization pool, and the number of neurons per area held fixed. The free parameters were the gains, the time constants, the noise parameters, the semi-saturation constants $\sigma_{1,2}$, and $\alpha_{1,2}$ (32 free parameters in total; Tables S1 and S3). The power and coherence spectra were fitted separately, giving two parameter sets. The objective, the mean squared error between the model and the experimental spectra over all contrasts and frequencies, was minimized with Adam on a bounded reparametrization of the parameters. Gradients were obtained by automatic differentiation through the fixed point (implicit differentiation), the linearization, and the spectra. The power fit used all eight contrasts with equal weight.

The fitted parameters remain close to the baseline set: $\gamma_1 \approx 1.07$ and $\sigma_1 \approx 0.077$ (baseline values 1.0 and 0.07), $\beta_1 \approx 0.7$, $\beta_2 \approx 1.2$–$1.4$, $\alpha_1 \approx 14$–$20$ (baseline 10), and all time constants between 0.2 and 3 ms. The resulting fits are shown in Fig. S7: the root-mean-square deviation is below 0.062 for the normalized power and below 0.048 for the coherence at every contrast.

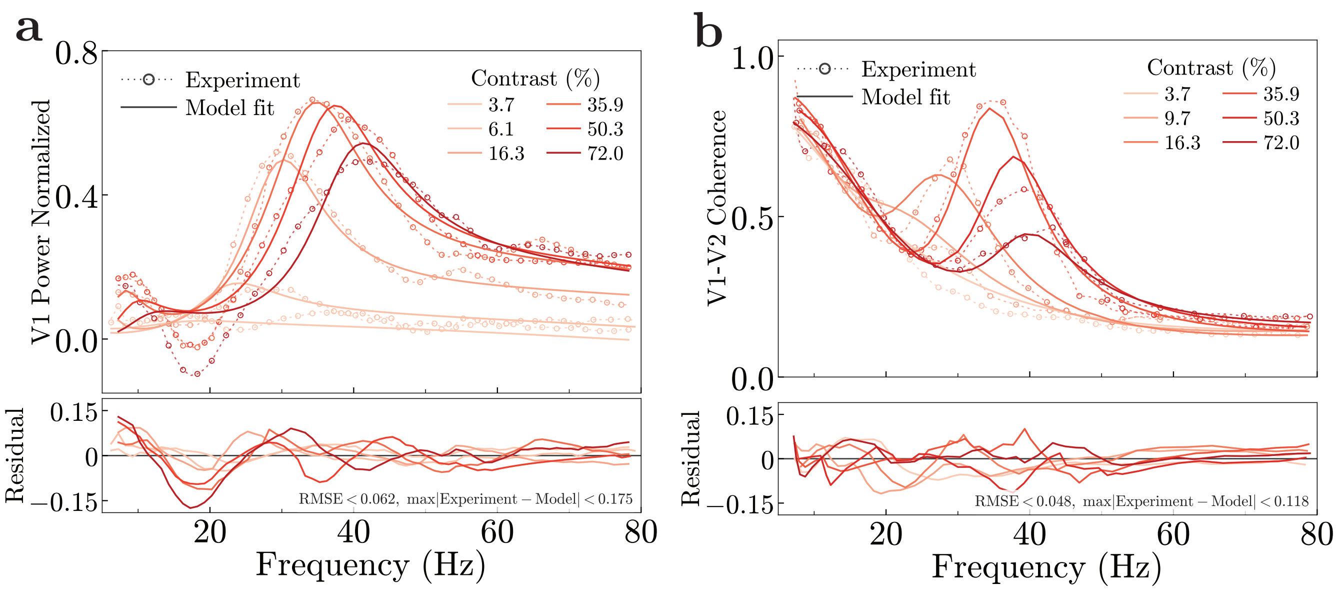


FIG. S7. **Quantitative fit of the theory to experimental power and coherence spectra.** Unlike Fig. 2 of the main text, in which the theory is compared with data using a single, unfitted set of baseline parameters, here the model parameters were optimized to fit the experimental spectra measured in macaque V1 and V2 [2]. Dotted lines with circles: experiment; solid lines: model fit. Line color indicates stimulus contrast (%; legend). **a,** Normalized V1 power spectra at six contrast levels (3.7%, 6.1%, 16.3%, 35.9%, 50.3%, and 72.0%). **b,** V1–V2 coherence spectra at six contrast levels (3.7%, 9.7%, 16.3%, 35.9%, 50.3%, and 72.0%). Bottom panels show the residual (Experiment − Model) at each contrast. The fitted model captures the shift of the gamma peak toward higher frequencies with increasing contrast, the non-monotonic dependence of peak amplitude on contrast, and the high coherence at low frequencies for low contrasts.

### IV. Robustness with respect to model parameters

All the results in this paper are obtained from the same set of parameters (Table S1) unless otherwise specified. To ensure the robustness of our findings, we verified them against a wide range of model parameters. We systematically

| **Parameter** | **Value** |
|---|---|
| $\tau_{\mathbf{y_1},\mathbf{u_1},\mathbf{a_1},\mathbf{q_1}}$ | 1 ms |
| $\tau_{\mathbf{y_2},\mathbf{u_2},\mathbf{a_2},\mathbf{q_2}}$ | 1 ms |
| $\gamma_1$ | 1.0 |
| $\beta_{\{1,2\}}$ | 1.0 |
| $\alpha_{\{1,2\}}$ | 10 |
| $\sigma_{\{1,2\}}$ | 0.07 |

TABLE S1. Default parameter values

| **Parameter** | **Stability Range** |
|---|---|
| $\gamma_1$ | [0.0 – 1.1] |
| $\beta_{\{1,2\}}$ | [0.0 – 2.5] |
| $\alpha_{\{1,2\}}$ | [7.5 – 50] |
| $\sigma_{\{1,2\}}$ | [0.035 – 0.14] |

TABLE S2. Valid parameter ranges

repeated the calculations and simulations described above, varying each parameter individually while holding others at their baseline values. These parameter ranges are specified in Table S2. Overall, our key findings remain robust across a wide range of parameter regimes without requiring precise parameter tuning:

- Contrast-response functions (Fig. 2a). Higher cortical area (V2) consistently exhibits greater contrast gain, reflected by a lower semi-saturation contrast and a steeper exponent than lower cortical area (V1).

- Gamma oscillations (Figs. 2c,e). Both the power spectra and coherence spectra consistently exhibit prominent gamma-band peaks that shift to higher frequencies as contrast increases.

- Alpha oscillations (Figs. 3c). Low contrast inputs and weak feedback gain consistently produce clear alpha-band peaks in the power spectra.

- Communication subspaces (Fig. 2g). The V1-V2 subspace is consistently lower dimensional than the V1-V1 subspace.
- Communication subspaces and feedback gain (Fig. 3k). The V1-V2 communication strength consistently increases with increasing feedback gain.
- Communication subspaces and V1 input gain (Fig. 3l). V1-V1 and V1-V2 communication strength consistently decrease with increasing V1 input gain.
- V2 input gain (Suppl. Fig. S8). Increasing the V2 input gain $\beta_2$ consistently enhances inter-areal (V1-V2) communication while leaving within-area (V1-V1) communication approximately unchanged, and shifts the gamma peaks in power and coherence to higher frequencies while broadening them and reducing their amplitude.
- Frequency-dependence of communication subspaces (Fig. 4). The communication strength peaks consistently at the same frequencies as coherence for each contrast level, and subspace dimensionality is minimized at those frequencies.
- Dynamic functional connectivity (Fig. 5). Enhancing feedback from V4 to V1, consistently boosts communication between V1 and V4, and likewise, enhancing feedback from V5 to V1, selectively enhances communication between V1 and V5.
- Dimensionality of inter-areal communication subspaces (Suppl. Figs. S4l,p). The dimensionality is consistently low when the input drive is localized (e.g., gratings). However, delocalized input drives (e.g., naturalistic images) result in a higher dimensionality. Fully all-to-all connections between V1 and V2 also produce a higher subspace dimensionality.
- Oscillatory activity depends on normalization. Removing normalization consistently eliminates the gamma band peaks in the power spectrum, coherence spectrum, and communication strength (Suppl. Figs. S4e,f,g).
- The dimensionality of the communication subspace is governed by the rank of the cross-covariance matrix, which itself depends crucially on normalization (Suppl. Fig. S5).

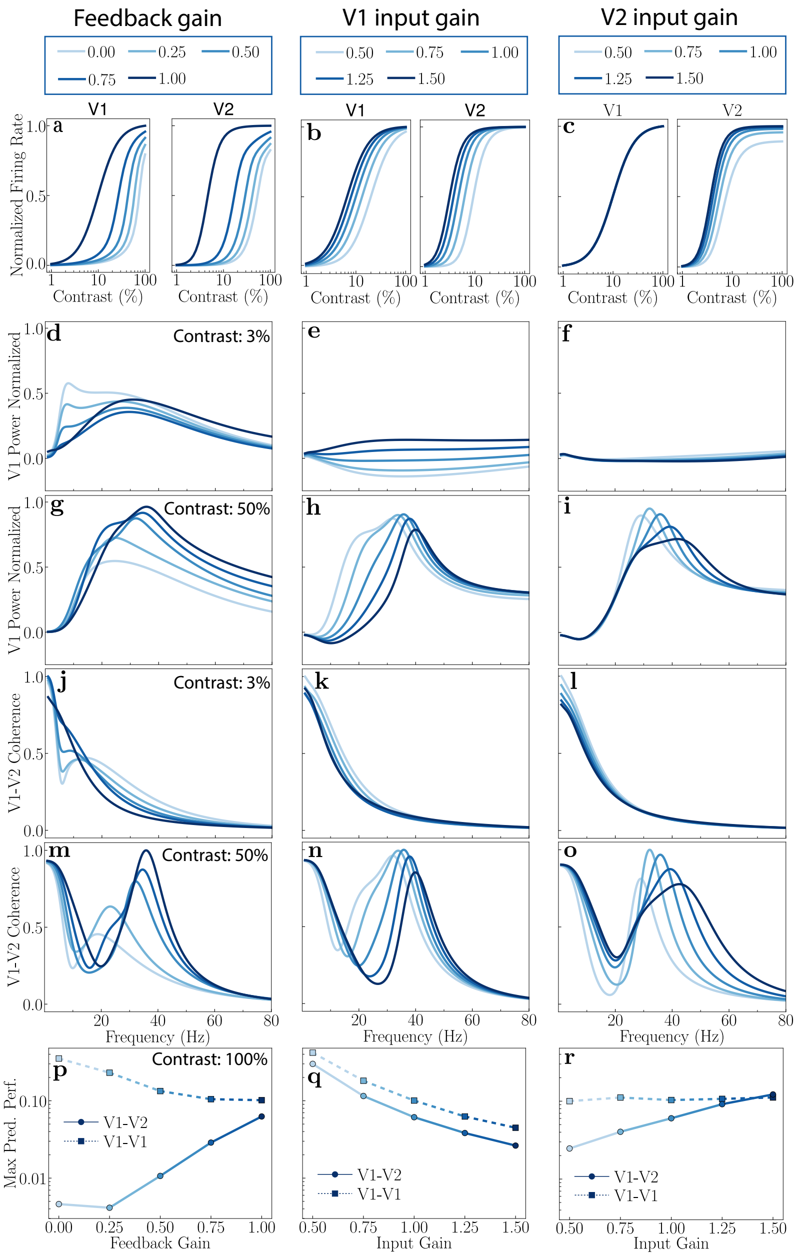


FIG. S8. **Theoretical predictions: Modulating feedback gain ($\gamma_1$; left column), V1 input gain ($\beta_1$; middle column), and V2 input gain ($\beta_2$; right column).** This figure extends Fig. 3 of the main text to include modulation of the V2 input gain (the gain of the V1 $\rightarrow$ V2 drive) on the same footing as feedback and V1 input gain. Line color intensity indicates the gain value (legends at top). Feedback gain was varied over $\gamma_1 \in \{0, 0.25, 0.5, 0.75, 1\}$ and input gains over $\beta_{1,2} \in \{0.5, 0.75, 1, 1.25, 1.5\}$, with the other two gains held at their baseline values. In **p–r**, source and target populations of 30 neurons were drawn at random from the 72 principal neurons of each area, and the prediction performance was averaged over 200 random draws. **a–c,** Firing rates as a function of contrast in V1 and V2. Increasing either feedback gain (**a**) or V1 input gain (**b**) enhances responses in both areas, whereas V2 input gain (**c**) leaves V1 responses unchanged and enhances only V2 responses. **d–f,** V1 power spectra at 3% contrast. The alpha peak observed at low feedback gain (**d**) is absent when V1 input gain (**e**) or V2 input gain (**f**) is varied. **g–i,** V1 power spectra at 50% contrast. The gamma peak shifts toward higher frequencies with increasing feedback gain (**g**), V1 input gain (**h**), and V2 input gain (**i**). Its bandwidth narrows with increasing V1 input gain but broadens with increasing feedback gain or V2 input gain, and its amplitude decreases with increasing V2 input gain. **j–l,** V1–V2 coherence spectra at 3% contrast. A broad beta-band peak is observed only at low feedback gain (**j**); no such peak is observed for changes in V1 input gain (**k**) or V2 input gain (**l**). **m–o,** V1–V2 coherence spectra at 50% contrast. The gamma coherence peak shifts toward higher frequencies with increasing feedback gain (**m**), V1 input gain (**n**), and V2 input gain (**o**); with increasing V2 input gain, the peak also broadens and its amplitude decreases. **p–r,** Maximum prediction performance of inter-areal (V1–V2, circles) and within-area (V1–V1, squares) communication subspaces at 100% contrast, as a function of each gain. Increasing feedback gain enhances inter-areal communication while reducing within-area communication (**p**); increasing V1 input gain reduces both (**q**); increasing V2 input gain enhances inter-areal communication while leaving within-area communication approximately unchanged (**r**).

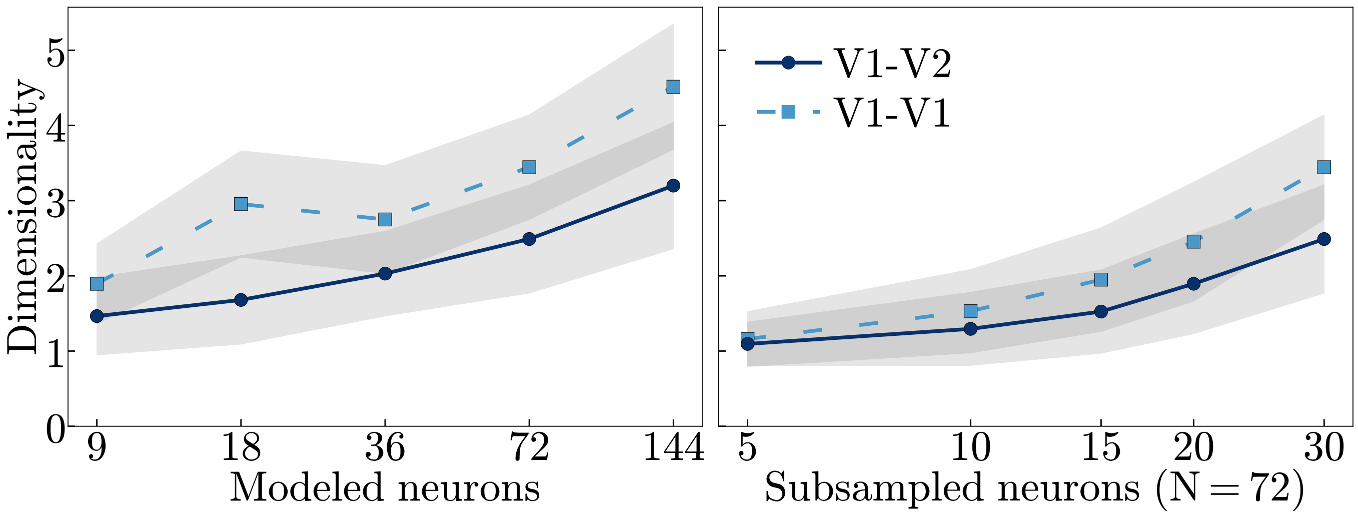


FIG. S9. **Dependence of communication-subspace dimensionality on the number of modeled and sampled neurons.** Dimensionality of the inter-areal (V1–V2, circles) and within-area (V1–V1, squares) communication subspaces at 100% contrast and baseline parameters, defined as the smallest number of predictive dimensions at which the prediction performance reaches 95% of its full-rank value. Symbols show the mean, and shaded regions ±1 standard deviation, over 200 random draws of source and target populations. **Left,** The number of modeled principal neurons per area was varied ($N$ = 9, 18, 36, 72, and 144) while the sampled fraction was held at a fixed value (≈ 42%). The connectivity kernels and the normalization pool are defined as functions of position on the ring and therefore scale with $N$; no other parameter was changed. **Right,** The number of sampled source and target neurons was varied (5–30) at the value $N$ = 72 used in the main text. In both cases the dimensionality grows slowly with the number of neurons (roughly two-fold over the 16-fold change in $N$) and remains far below the number of sampled neurons, and the inter-areal subspace is lower-dimensional than the within-area subspace at every population size (the two coincide only for the smallest samples, where both approach one). Absolute dimensionality values therefore depend on the population size, and comparisons with experimental estimates should be made at matched population sizes, as in Fig. 2g,h of the main text.

### V. Derivation of the General Hierarchical Recurrent Normalization Circuit

This appendix provides a detailed derivation of the general dynamical equations for the hierarchical recurrent normalization model with reciprocal connections. We begin with the primary visual cortex (V1). Assuming $\mathbf{N}_1$ is the normalization weight matrix for V1 and $\mathbf{z}_1$ is the input drive to V1, we can write the normalization equations for the V1 principal neurons with complementary receptive fields as:

$$\mathbf{y}_1^{s+} = \frac{\lfloor \mathbf{z}_1 \rfloor^n}{\boldsymbol{\sigma}^n + \mathbf{N}_1(\lfloor \mathbf{z}_1 \rfloor^n + \lfloor -\mathbf{z}_1 \rfloor^n)} \qquad \mathbf{y}_1^{s-} = \frac{\lfloor -\mathbf{z}_1 \rfloor^n}{\boldsymbol{\sigma}^n + \mathbf{N}_1(\lfloor \mathbf{z}_1 \rfloor^n + \lfloor -\mathbf{z}_1 \rfloor^n)} \tag{S1}$$

Where $\mathbf{y}_1^{s+}$ and $\mathbf{y}_1^{s-}$ represent the steady-state firing rates of principal neurons corresponding to positive ($\lfloor \mathbf{z}_1 \rfloor^n$) and negative $\lfloor -\mathbf{z}_1 \rfloor^n$ input drives, respectively. Here, $\sigma$ represents the semi-saturation constant. The exponent $n$ is typically observed to be $n \approx 2$ for the primary visual cortex, it can be different for different brain areas. Mathematically:

$$\lfloor \mathbf{z}_1 \rfloor^n + \lfloor -\mathbf{z}_1 \rfloor^n = |\mathbf{z}_1|^n \tag{S2}$$

Our goal is to derive the dynamical equations for a general $n$, that yield the fixed points defined by Eq. S1.
First, we distinguish between the membrane potentials and the firing rates of neurons. The membrane potential of a given type of neuron is denoted by vectors such as $\mathbf{y}_1$, $\mathbf{u}_1$, $\mathbf{a}_1$, and $\mathbf{q}_1$. Their corresponding firing rates are $\mathbf{y}_1^+$ (and $\mathbf{y}_1^-$ for complementary responses), $\mathbf{u}_1^+$, $\mathbf{a}_1^+$, and $\mathbf{q}_1^+$. The instantaneous firing rates are related to membrane potentials by applying rectification (denoted by $\lfloor \cdot \rfloor$) followed by power-law activation [3],

$$\mathbf{x}^+ = \lfloor \mathbf{x} \rfloor^\beta. \tag{S3}$$

The values of $\beta$ are taken to be $\{2,\frac{1}{2},1,1\}$ for $\{\mathbf{y}_1,\ \mathbf{u}_1,\ \mathbf{a}_1,\ \mathbf{q}_1\}$, respectively. Combining the firing rates of principal neurons with the complementary receptive fields, $\mathbf{y}_1^{s+}$ and $\mathbf{y}_1^{s-}$, Eq. S1 can be alternatively written as,

$$|\mathbf{y}_1|^n = \mathbf{y}_1^{s+} + \mathbf{y}_1^{s-} = \frac{|\mathbf{z}_1|^n}{\boldsymbol{\sigma}^n + \mathbf{N}_1 |\mathbf{z}_1|^n} \tag{S4}$$

To proceed with our derivation, we need to isolate the $|\mathbf{z}_1|^n$ term. From Eq. S4, we have:

$$|\mathbf{z}_1|^n = \boldsymbol{\sigma}^n \odot |\mathbf{y}_1|^n + |\mathbf{y}_1|^n \odot (\mathbf{N}_1 |\mathbf{z}_1|^n) \tag{S5}$$

where $\odot$ denotes element-wise multiplication. Using the identity $\mathbf{x}_1 \odot \mathbf{x}_2 = \mathrm{D}(\mathbf{x}_1)\mathbf{x}_2$, where $\mathrm{D}(\mathbf{x}_1)$ is a diagonal matrix with the elements of vector $\mathbf{x}_1$ on its diagonal, we can rewrite and solve for $|\mathbf{z}_1|^n$:

$$|\mathbf{z}_1|^n = \boldsymbol{\sigma}^n \odot |\mathbf{y}_1|^n + \mathrm{D}(|\mathbf{y}_1|^n)\mathbf{N}_1 |\mathbf{z}_1|^n \tag{S6}$$

Factoring out $|\mathbf{z}_1|^n$:

$$[\mathbf{I} - \mathrm{D}(|\mathbf{y}_1|^n)\,\mathbf{N}_1]\,|\mathbf{z}_1|^n = \boldsymbol{\sigma}^n \odot |\mathbf{y}_1|^n \tag{S7}$$

Multiplying both sides with $[\mathbf{I} - \mathrm{D}(|\mathbf{y}_1|^n)\mathbf{N}_1]^{-1}$ from the left yields:

$$|\mathbf{z}_1|^n = [\mathbf{I} - \mathrm{D}(|\mathbf{y}_1|^n)\mathbf{N}_1]^{-1} (\boldsymbol{\sigma}^n \odot |\mathbf{y}_1|^n) \tag{S8}$$

This expression for $|\mathbf{z}_1|^n$ will be used in subsequent steps. Now, we introduce the dynamical equation for the membrane potential $\mathbf{y}_1$ of the principal V1 neurons, consistent with biological computation in pyramidal cells (see main text, "Biological plausibility"):

$$\tau_{y_1}\frac{\mathrm{d}\mathbf{y}_1}{\mathrm{d}t} = -\mathbf{y}_1 + \mathbf{b}_1 \odot \mathbf{z}_1 + \frac{1}{1+\mathbf{a}_1^+} \odot \left(\mathbf{W}_{11}\left((\mathbf{y}_1^+)^{\frac{1}{n}} - (\mathbf{y}_1^-)^{\frac{1}{n}}\right) + \boldsymbol{\gamma}_1 \odot \mathbf{W}_{12}(\mathbf{y}_2^+)^{\frac{1}{n}}\right). \tag{S9}$$

Here, $\tau_{y_1}$ is the time constant, $\mathbf{b}_1$ is the input gain, $\mathbf{a}_1^+$ represents firing rates of the inhibitory neuron population. $\mathbf{W}_{11}$ is the recurrent V1 connectivity matrix, $\mathbf{W}_{12}$ is the connectivity matrix containing the synaptic weights from V2 to V1. $\mathbf{W}_{12}(\mathbf{y}_2^+)^{\frac{1}{n}}$ represents the feedback signal from V2 to V1, calculated as the weighted sum of the activities of V2 neurons. $\boldsymbol{\gamma}_1$ is the feedback gain modulation parameter. $\mathbf{y}_2^+$ is the firing rate of V2 principal neurons, defined as $\lfloor \mathbf{y}_2 \rfloor^n$.

For simplicity, we assume that the recurrent weight matrix $\mathbf{W}_{11} = \mathrm{I}$. Given that $(\mathbf{y}_1^+)^{1/n} - (\mathbf{y}_1^-)^{1/n} = \lfloor \mathbf{y}_1 \rfloor - \lfloor -\mathbf{y}_1 \rfloor = \mathbf{y}_1$ and $(\mathbf{y}_2^+)^{1/n} = \lfloor \mathbf{y}_2 \rfloor$, the equation for $\mathbf{y}_1$ simplifies to:

$$\tau_{y_1}\frac{\mathrm{d}\mathbf{y}_1}{\mathrm{d}t} = -\mathbf{y}_1 + \mathbf{b}_1 \odot \mathbf{z}_1 + \frac{1}{1+\mathbf{a}_1^+} \odot (\mathbf{y}_1 + \boldsymbol{\gamma}_1 \odot \mathbf{W}_{12}\lfloor \mathbf{y}_2 \rfloor) \tag{S10}$$

Now at steady state, $\frac{\mathrm{d}\mathbf{y}_1}{\mathrm{d}t} = 0$,

$$0 = -\mathbf{y}_1 + \mathbf{b}_1 \odot \mathbf{z}_1 + \frac{1}{1+\mathbf{a}_1^+} \odot (\mathbf{y}_1 + \boldsymbol{\gamma}_1 \odot \mathbf{W}_{12}\lfloor \mathbf{y}_2 \rfloor) \tag{S11}$$

Expanding the terms:

$$0 = -\mathbf{y}_1 + \mathbf{b}_1 \odot \mathbf{z}_1 + \frac{1}{1+\mathbf{a}_1^+} \odot \mathbf{y}_1 + \frac{1}{1+\mathbf{a}_1^+} \odot (\boldsymbol{\gamma}_1 \odot \mathbf{W}_{12}\lfloor \mathbf{y}_2 \rfloor) \tag{S12}$$

Rearranging terms:

$$\frac{\mathbf{a}_1^+}{1+\mathbf{a}_1^+} \odot \mathbf{y}_1 = \mathbf{b}_1 \odot \mathbf{z}_1 + \frac{1}{1+\mathbf{a}_1^+} \odot (\boldsymbol{\gamma}_1 \odot \mathbf{W}_{12}\lfloor \mathbf{y}_2 \rfloor) \tag{S13}$$

Moving the feedback term to the left-hand side:

$$\frac{\mathbf{a}_1^+}{1+\mathbf{a}_1^+} \odot \mathbf{y}_1 - \frac{1}{1+\mathbf{a}_1^+} \odot (\boldsymbol{\gamma}_1 \odot \mathbf{W}_{12}\lfloor \mathbf{y}_2 \rfloor) = \mathbf{b}_1 \odot \mathbf{z}_1 \tag{S14}$$

Factoring out $\mathbf{y}_1$:

$$\mathbf{y}_1 \odot \left( \frac{\mathbf{a}_1^+}{1+\mathbf{a}_1^+} - \frac{1}{1+\mathbf{a}_1^+} \odot \frac{\boldsymbol{\gamma}_1 \odot \mathbf{W}_{12}\lfloor \mathbf{y}_2 \rfloor}{\mathbf{y}_1} \right) = \mathbf{b}_1 \odot \mathbf{z}_1 \tag{S15}$$

We define a modulatory term $\mathbf{u}_1^+$, such that:

$$\mathbf{y}_1 \odot \mathbf{u}_1^+ = \mathbf{b}_1 \odot \mathbf{z}_1 \tag{S16}$$

where:

$$\mathbf{u}_1^+ = \frac{\mathbf{a}_1^+}{1+\mathbf{a}_1^+} - \frac{1}{1+\mathbf{a}_1^+} \odot \frac{\boldsymbol{\gamma}_1 \odot \mathbf{W}_{12}\lfloor \mathbf{y}_2 \rfloor}{\mathbf{y}_1} \tag{S17}$$

Next, we derive the dynamical equation for $\mathbf{a}_1$, from the definition of $\mathbf{u}_1^+$:

$$\mathbf{u}_1^+ \odot (1+\mathbf{a}_1^+) = \mathbf{a}_1^+ - \frac{\boldsymbol{\gamma}_1 \odot \mathbf{W}_{12}\lfloor \mathbf{y}_2 \rfloor}{\mathbf{y}_1} \tag{S18}$$

Upon rearranging,

$$0 = -\mathbf{a}_1^+ + \frac{\boldsymbol{\gamma}_1 \odot \mathbf{W}_{12}\lfloor \mathbf{y}_2 \rfloor}{\mathbf{y}_1} + (1+\mathbf{a}_1^+) \odot \mathbf{u}_1^+ \tag{S19}$$

This equation describes the steady state of $\mathbf{a}_1$. It exhibits a form analogous to that of $\mathbf{y}_1$, comprising an input drive and a recurrent drive term. Therefore, we propose the following dynamical equation for the membrane potential $\mathbf{a}_1$:

$$\tau_{a_1} \frac{\mathrm{d}\mathbf{a}_1}{\mathrm{d}t} = -\mathbf{a}_1 + \frac{\boldsymbol{\gamma}_1 \odot \mathbf{W}_{12}\lfloor \mathbf{y}_2 \rfloor}{\mathbf{y}_1} + (1+\mathbf{a}_1^+) \odot \mathbf{u}_1^+ \tag{S20}$$

According to Dale's Law, the principal neurons $\mathbf{y}_1$, which we hypothesize to be excitatory (main text, "Biological plausibility"), cannot directly inhibit other neurons. Therefore, the division by $\mathbf{y}_1$ implies the existence of an intermediate inhibitory interneuron. We introduce an inhibitory interneuron population $\mathbf{q}_1$, that tracks $\mathbf{y}_1$:

$$\tau_{q_1} \frac{\mathrm{d}\mathbf{q}_1}{\mathrm{d}t} = -\mathbf{q}_1 + (\mathbf{y}_1^+)^{1/n} \tag{S21}$$

With this interneuron, the dynamic equation for $\mathbf{a}_1$ becomes:

$$\tau_{a_1} \frac{\mathrm{d}\mathbf{a}_1}{\mathrm{d}t} = -\mathbf{a}_1 + \frac{\boldsymbol{\gamma}_1 \odot \mathbf{W}_{12}(\mathbf{y}_2^+)^{1/n}}{\mathbf{q}_1^+} + (1+\mathbf{a}_1^+) \odot \mathbf{u}_1^+ \tag{S22}$$

Now, we return to Eq. S16 to derive the dynamical equation for the modulatory neuron $\mathbf{u}_1$. Starting with $\mathbf{y}_1 \odot \mathbf{u}_1^+ = \mathbf{b}_1 \odot \mathbf{z}_1$, we take the modulus on both sides and raise to the power of n:

$$|\mathbf{y}_1|^n \odot (\mathbf{u}_1^+)^n = \mathbf{b}_1^n \odot |\mathbf{z}_1|^n \tag{S23}$$

Substituting the expression for $|\mathbf{z}_1|^n$ from Eq. S8:

$$|\mathbf{y}_1|^n \odot (\mathbf{u}_1^+)^n = \mathbf{b}_1^n \odot [\mathbf{I} - \mathrm{D}(|\mathbf{y}_1|^n)\mathbf{N}_1]^{-1}(\boldsymbol{\sigma}^n \odot |\mathbf{y}_1|^n) \tag{S24}$$

We assume that $\mathbf{b}_1$ has equal components, i.e., $\mathbf{b}_1 = b_1\mathbf{1}$. Multiplying both sides by $[\mathbf{I} - \mathrm{D}(|\mathbf{y}_1|^n)\mathbf{N}_1]$ and rearranging:

$$[\mathbf{I} - \mathrm{D}(|\mathbf{y}_1|^n)\mathbf{N}_1] \left( \frac{(\mathbf{u}_1^+)^n \odot |\mathbf{y}_1|^n}{\mathbf{b}_1^n} \right) = \boldsymbol{\sigma}^n \odot |\mathbf{y}_1|^n \tag{S25}$$

Multiplying both sides by $\mathrm{D}\left(\frac{1}{|\mathbf{y}_1|^n}\right)$:

$$\mathrm{D}\left(\frac{1}{|\mathbf{y}_1|^n}\right)[\mathbf{I}-\mathrm{D}(|\mathbf{y}_1|^n)\mathbf{N}_1]\left(\frac{(\mathbf{u}_1^+)^n\odot|\mathbf{y}_1|^n}{\mathbf{b}_1^n}\right)=\mathrm{D}\left(\frac{1}{|\mathbf{y}_1|^n}\right)(\boldsymbol{\sigma}^n\odot|\mathbf{y}_1|^n) \tag{S26}$$

This simplifies to:

$$\frac{(\mathbf{u}_1^+)^n}{\mathbf{b}_1^n}-\mathbf{N}_1\left(\frac{(\mathbf{u}_1^+)^n\odot|\mathbf{y}_1|^n}{\mathbf{b}_1^n}\right)=\boldsymbol{\sigma}^n \tag{S27}$$

Since we assumed that $\mathbf{b}_1$ has equal components, we get:

$$(\mathbf{u}_1^+)^n-\mathbf{N}_1((\mathbf{u}_1^+)^n\odot|\mathbf{y}_1|^n)=\boldsymbol{\sigma}^n\odot\mathbf{b}_1^n \tag{S28}$$

Elementwise multiplying both sides by $\mathbf{u}_1/(\mathbf{u}_1^+)^n$ from left and rearrange:

$$0=-\mathbf{u}_1+\frac{\mathbf{u}_1}{(\mathbf{u}_1^+)^n}\odot\left(\mathbf{b}_1^n\odot\boldsymbol{\sigma}^n+\mathbf{N}_1((\mathbf{u}_1^+)^n\odot|\mathbf{y}_1|^n)\right) \tag{S29}$$

This equation describes the steady state of $\mathbf{u}_1$. We propose the following dynamical equation for $\mathbf{u}_1$,

$$\tau_{u_1}\frac{\mathrm{d}\mathbf{u}_1}{\mathrm{d}t}=-\mathbf{u}_1+\frac{\mathbf{u}_1}{(\mathbf{u}_1^+)^n}\odot\left(\mathbf{b}_1^n\odot\boldsymbol{\sigma}^n+\mathbf{N}_1((\mathbf{u}_1^+)^n\odot|\mathbf{y}_1|^n)\right) \tag{S30}$$

These equations (Eq. S10, Eq. S30, Eq. S22, Eq. S21) describe the dynamics of principal neurons, modulatory inhibitory neurons, and modulatory excitatory neurons in V1. They exhibit the fixed point solution given by the normalization by design. These equations are general for any choice of $n$ and $\beta$. We get a particularly simplified form when we set $n=2$ and $\beta=\{2,\frac{1}{2},1,1\}$ for $\{\mathbf{y}_1,\,\mathbf{u}_1,\,\mathbf{a}_1,\,\mathbf{q}_1\}$, respectively.
Finally we rewrite the set of four dynamical equations for V1 as:

$$\tau_{y_1}\frac{\mathrm{d}\mathbf{y}_1}{\mathrm{d}t}=-\mathbf{y}_1+\mathbf{b}_1^{(y_1)}\odot\mathbf{z}_1+\left(\frac{1}{1+\mathbf{a}_1^+}\right)\left(\mathbf{W}_{11}\sqrt{\mathbf{y}_1^+}+\boldsymbol{\gamma}_1^{(y_1)}\odot\mathbf{W}_{12}\sqrt{\mathbf{y}_2^+}\right) \tag{S31}$$

$$\tau_{u_1}\frac{\mathrm{d}\mathbf{u}_1}{\mathrm{d}t}=-\mathbf{u}_1+((\mathbf{b}_1^{(u_1)})^2\odot\boldsymbol{\sigma}^2)+\mathbf{N}_1\left(\mathbf{y}_1^+\odot(\mathbf{u}_1^+)^2\right) \tag{S32}$$

$$\tau_{a_1}\frac{\mathrm{d}\mathbf{a}_1}{\mathrm{d}t}=-\mathbf{a}_1+\frac{\boldsymbol{\gamma}_1^{(a_1)}\odot\mathbf{W}_{12}\sqrt{\mathbf{y}_2^+}}{\mathbf{q}_1^+}+\mathbf{u}_1^++\mathbf{a}_1^+\odot\mathbf{u}_1^++\alpha_1\frac{\mathrm{d}\mathbf{u}_1}{\mathrm{d}t} \tag{S33}$$

$$\tau_{q_1}\frac{\mathrm{d}\mathbf{q}_1}{\mathrm{d}t}=-\mathbf{q}_1+\sqrt{\mathbf{y}_1^+} \tag{S34}$$

where we have decoupled the input gain $\mathbf{b}_1$ and feedback gain $\boldsymbol{\gamma}_1$ across each equation. When $\mathbf{b}_1^{(y_1)}=\mathbf{b}_1^{(u_1)}$ and $\boldsymbol{\gamma}_1^{(y_1)}=\boldsymbol{\gamma}_1^{(a_1)}$, and $\mathbf{W}_{11}=\mathbf{I}$ we exactly recover the normalization equation, Eq. 2, at steady state for $\mathbf{y}_1$. Then to reduce the number of parameters we re-define the input gain as $\boldsymbol{\beta}_1\equiv\mathbf{b}_1^{(y_1)}/\mathbf{b}_1^{(u_1)}$ and fix $\mathbf{b}_1^{(u_1)}=1/2$. Similarly, we re-define the feedback gain as $\boldsymbol{\gamma}_1\equiv\boldsymbol{\gamma}_1^{(y_1)}/\boldsymbol{\gamma}_1^{(a_1)}$ and fix $\boldsymbol{\gamma}_1^{(a_1)}=1/2$. We thus arrive at Eq. 1 of the main text. Finally, note that the term $\alpha_1\frac{\mathrm{d}\mathbf{u}_1}{\mathrm{d}t}$ is added to the dynamics of $\mathbf{a}_1$ to achieve correct oscillatory behavior. This term does not impact the steady state solution but influences the dynamics.

The dynamical equations for V2 can be derived following a similar procedure:

$$\tau_{y_2}\frac{\mathrm{d}\mathbf{y}_2}{\mathrm{d}t}=-\mathbf{y}_2+\mathbf{b}_2^{(y_2)}\odot\mathbf{z}_2+\left(\frac{1}{1+\mathbf{a}_2^+}\right)\left(\mathbf{W}_{22}\sqrt{\mathbf{y}_2^+}\right) \tag{S35}$$

$$\tau_{u_2}\frac{\mathrm{d}\mathbf{u}_2}{\mathrm{d}t}=-\mathbf{u}_2+((\mathbf{b}_2^{(u_2)})^2\odot\boldsymbol{\sigma}^2)+\mathbf{N}_2\left(\mathbf{y}_2^+\odot(\mathbf{u}_2^+)^2\right) \tag{S36}$$

$$\tau_{a_2}\frac{\mathrm{d}\mathbf{a}_2}{\mathrm{d}t}=-\mathbf{a}_2+\mathbf{u}_2^++\mathbf{a}_2^+\odot\mathbf{u}_2^++\alpha_2\frac{\mathrm{d}\mathbf{u}_2}{\mathrm{d}t} \tag{S37}$$

$$\tau_{q_2}\frac{\mathrm{d}\mathbf{q}_2}{\mathrm{d}t}=-\mathbf{q}_2+\sqrt{\mathbf{y}_2^+} \tag{S38}$$

where, $\mathbf{z}_2 = \mathbf{W}_{21}\mathbf{y}_1^+$; and, like for $\mathbf{y}_1$, we have decoupled the input gain $\mathbf{b}_2$ across each equation. When $\mathbf{b}_2^{(y_2)} = \mathbf{b}_2^{(u_2)}$ and $\mathbf{W}_{22} = \mathbf{I}$ we exactly recover the normalization equation, Eq. 2, at steady state for $\mathbf{y}_2$. To reduce the number of parameters, we redefine the input gain as $\boldsymbol{\beta}_2 \equiv \mathbf{b}_2^{(y_2)}/\mathbf{b}_2^{(u_2)}$ and fix $\mathbf{b}_2^{(u_2)} = \mathbf{b}_2^{(y_2)} = 1/2$. Note that in principle $\mathbf{y}_2$ can receive feedback from higher cortical areas, in which case the equations would look identical to Eq. 1 but the subscripts change as $1 \rightarrow 2$ and $2 \rightarrow 3$.

In addition to the membrane potential, the corresponding firing rates of V1 and V2 neurons are also simulated. These are modeled as low-pass filtered versions of their target rectified power-law transformations. For V1, as an example:

$$\tau_{y_1^+}\frac{\mathrm{d}\mathbf{y}_1^+}{\mathrm{d}t} = -\mathbf{y}_1^+ + \lfloor \mathbf{y}_1 \rfloor^2 \tag{S39}$$

$$\tau_{u_1^+}\frac{\mathrm{d}\mathbf{u}_1^+}{\mathrm{d}t} = -\mathbf{u}_1^+ + \sqrt{\lfloor \mathbf{u}_1 \rfloor} \tag{S40}$$

$$\tau_{a_1^+}\frac{\mathrm{d}\mathbf{a}_1^+}{\mathrm{d}t} = -\mathbf{a}_1^+ + \lfloor \mathbf{a}_1 \rfloor \tag{S41}$$

$$\tau_{q_1^+}\frac{\mathrm{d}\mathbf{q}_1^+}{\mathrm{d}t} = -\mathbf{q}_1^+ + \lfloor \mathbf{q}_1 \rfloor \tag{S42}$$

### VI. Noise Model

#### A. Intrinsic neural circuit noise

The influence of noise on the deterministic neural dynamics is treated analytically in this model. We consider two primary sources of noise: firing rate noise originating from the spiking process and synaptic noise.

**i)** Firing rate noise is added to the firing rate vector $\mathbf{k}^+$, where $\mathbf{k}^+$ represents various neural population firing rates (e.g., excitatory like $\mathbf{y}_1^+, \mathbf{u}_1^+$or inhibitory like $\mathbf{a}_1^+, \mathbf{q}_1^+$). This noise is attributed to the spiking mechanism, whose strength is found by the modified Gaussian Rectification (GR) model [4, 5]. Due to the high degree of redundancy in neural circuits, the strength of this noise is quenched due to the averaging of these redundant signals. The result is that the noise has a negligible impact on the firing rate dynamics [6, 7]. For a given firing rate vector $\mathbf{k}^+$, with an associated activation exponent $\beta$, the noise dynamics are described by:

$$\tau_{k^+}\frac{\mathrm{d}\mathbf{k}^+}{\mathrm{d}t} = -\mathbf{k}^+ + \lfloor \mathbf{k} \rfloor^\beta + \sigma_k \cdot \boldsymbol{\eta}(t), \tag{S43}$$

where $\tau_{k^+}$ is the time constant for the firing rate dynamics of population $\mathbf{k}^+$, $\sigma_k$ represents the noise strengths for different neuron types, and $\boldsymbol{\eta}(t)$ is a vector of independent Gaussian white noise process, each with zero mean and unit variance.

**ii)** Synaptic noise is modeled as exponentially low-pass filtered Gaussian white noise ($\mathbf{f}$), given by the following SDE:

$$\tau_f\frac{\mathrm{d}\mathbf{f}}{\mathrm{d}t} = -\mathbf{f} + \sigma_f \cdot \boldsymbol{\eta}(t), \tag{S44}$$

where $\tau_f$ is the noise filtering time constant ($\tau_f = 1$ msec, approximating the time constant of synaptic transmission), and $\sigma_f$ represents the noise strength. This filtered noise ($\mathbf{f}$) is then added to the dynamical equations for the membrane potentials of each neuron, Eq. S31-S38, and it is the main source of intrinsic noise.

Upon incorporating these sources of noise, we get a high-dimensional nonlinear stochastic dynamical system with the following variables:

- Variables representing the membrane potential for different neuron types implementing normalization (Eq. S31-S38).
- Firing rates (Eq. S43) for each membrane potential variable.
- Filtered synaptic noise (Eq. S44) for each membrane potential variable.

We then linearize this dynamical system about the fixed point found by simulating the deterministic part of the dynamical system, which gives us the Jacobian matrix $\mathbf{J}$. We also construct the dispersion matrix $\mathbf{L}$ and the diffusion matrix $\mathbf{D}$ using the strength of the noise added in various dynamical variables. The definition of these matrices allows us to compute the covariance matrix and power spectra analytically as discussed in the Methods section of the main text.

#### B. Extrinsic noise in the local field potential

In addition to the noise sources detailed above, we also account for additional sources of noise that contribute to LFP power and coherence spectra. We consider two components of extrinsic noise: **i)** Correlated noise ($\boldsymbol{\zeta}_{\text{correlated}}$), assumed to stem from the volume conduction of electric fields generated by transmembrane current through the extracellular brain tissue, with negligible impact on spiking activity and firing rates [8–10]. **ii)** Uncorrelated noise ($\boldsymbol{\zeta}_{\text{uncorrelated}}$), that encompasses noise from various other sources including motion artifact, instrumentation noise, thermal noise, and biological sources of noise [11–13]. This extrinsic noise is added to the simulated neural responses (membrane potentials for different neuron types, denoted collectively by $\mathbf{x}$):

$$\mathbf{x}'(t) = \mathbf{x}(t) + \boldsymbol{\zeta}_{\text{uncorrelated}}(t) + \boldsymbol{\zeta}_{\text{correlated}}(t) \tag{S45}$$

Since these noise sources are uncorrelated with the responses of the neurons, the power spectrum of the measured signals $\boldsymbol{\mathcal{S}}'(\omega)$ is the sum of the spectrum of the neurons' activity $\boldsymbol{\mathcal{S}}(\omega)$ and the spectra of these noise components:

$$\boldsymbol{\mathcal{S}}'(\omega) = \boldsymbol{\mathcal{S}}(\omega) + \boldsymbol{\mathcal{S}}_{\text{uncorrelated}}(\omega) + \boldsymbol{\mathcal{S}}_{\text{correlated}}(\omega) \tag{S46}$$

The correlated noise is modeled to produce a $\frac{1}{\omega^4}$ power law scaling, which is achieved by doubly low-pass filtered white noise. Its power spectrum for each element is given by:

$$\boldsymbol{\mathcal{S}}_{\text{correlated, element}}(\omega) = \frac{\Sigma_{\text{corr}}}{(1+\tau_{\text{noise}}^2\omega^2)^2} \tag{S47}$$

The uncorrelated power spectrum $\boldsymbol{\mathcal{S}}_{\text{uncorrelated}}(\omega)$ is diagonal, with each diagonal term having a similar filtered form:

$$\boldsymbol{\mathcal{S}}_{\text{uncorrelated}}(\omega) = \mathrm{D}\left(\frac{\Sigma_{\text{uncorr}}}{(1+\tau_{\text{noise}}^2\omega^2)^2}\right) \tag{S48}$$

In our simulation, the ratio of power spectral densities of the correlated noise to the uncorrelated noise (i.e, $\Sigma_{\text{corr}}/\Sigma_{\text{uncorr}}$ for corresponding element) is set to 10 for all the results. The noise parameters used in the simulation are as follows,

| Noise Parameter | Value |
|---|---|
| $\tau_x$ | 1 ms |
| $\tau_f$ | 1 ms |
| $\tau_{\text{noise}}$ | 50 ms |
| $\sigma_f$ | 0.01 |
| $\Sigma_{\text{corr}}$ | $9 \times 10^{-4}$ |
| $\Sigma_{\text{uncorr}}$ | $9 \times 10^{-5}$ |

TABLE S3. Noise parameters used in the simulation.


[1] J. D. Semedo, A. Zandvakili, C. K. Machens, M. Y. Byron, and A. Kohn, Cortical areas interact through a communication subspace, Neuron **102**, 249 (2019).

[2] M. J. Roberts, E. Lowet, N. M. Brunet, M. Ter Wal, P. Tiesinga, P. Fries, and P. De Weerd, Robust gamma coherence between macaque v1 and v2 by dynamic frequency matching, Neuron **78**, 523 (2013).

[3] D. J. Heeger, Half-squaring in responses of cat striate cells, Visual neuroscience **9**, 427 (1992).

[4] S. Rawat, D. Heeger, and S. Martiniani, A comprehensive large-scale model of the primary visual cortex (v1), Bulletin of the American Physical Society (2024).

[5] M. Carandini, Amplification of trial-to-trial response variability by neurons in visual cortex, PLoS biology **2**, e264 (2004).
[6] C. Allen and C. F. Stevens, An evaluation of causes for unreliability of synaptic transmission., Proceedings of the National Academy of Sciences **91**, 10380 (1994).
[7] Z. F. Mainen and T. J. Sejnowski, Reliability of spike timing in neocortical neurons, Science **268**, 1503 (1995).
[8] G. Buzsáki, C. A. Anastassiou, and C. Koch, The origin of extracellular fields and currents—eeg, ecog, lfp and spikes, Nature reviews neuroscience **13**, 407 (2012).
[9] C. Bédard, H. Kröger, and A. Destexhe, Model of low-pass filtering of local field potentials in brain tissue, Physical Review E—Statistical, Nonlinear, and Soft Matter Physics **73**, 051911 (2006).
[10] G. T. Einevoll, C. Kayser, N. K. Logothetis, and S. Panzeri, Modelling and analysis of local field potentials for studying the function of cortical circuits, Nature Reviews Neuroscience **14**, 770 (2013).
[11] M. J. Nelson and P. Pouget, Do electrode properties create a problem in interpreting local field potential recordings? (2010).
[12] K. A. Ludwig, R. M. Miriani, N. B. Langhals, M. D. Joseph, D. J. Anderson, and D. R. Kipke, Using a common average reference to improve cortical neuron recordings from microelectrode arrays, Journal of neurophysiology **101**, 1679 (2009).
[13] N. W. Whitmore and S.-C. Lin, Unmasking local activity within local field potentials (lfps) by removing distal electrical signals using independent component analysis, Neuroimage **132**, 79 (2016).
